# CRISPRmap: Sequencing-free optical pooled screens mapping multi-omic phenotypes in cells and tissue

**DOI:** 10.1101/2023.12.26.572587

**Authors:** Jiacheng Gu, Abhishek Iyer, Ben Wesley, Angelo Taglialatela, Giuseppe Leuzzi, Sho Hangai, Aubrianna Decker, Ruoyu Gu, Naomi Klickstein, Yuanlong Shuai, Kristina Jankovic, Lucy Parker-Burns, Yinuo Jin, Jia Yi Zhang, Justin Hong, Steve Niu, Jacqueline Chou, Dan A. Landau, Elham Azizi, Edmond M. Chan, Alberto Ciccia, Jellert T. Gaublomme

## Abstract

Pooled genetic screens are powerful tools to study gene function in a high-throughput manner. Typically, sequencing-based screens require cell lysis, which limits the examination of critical phenotypes such as cell morphology, protein subcellular localization, and cell-cell/tissue interactions. In contrast, emerging optical pooled screening methods enable the investigation of these spatial phenotypes in response to targeted CRISPR perturbations. In this study, we report a multi-omic optical pooled CRISPR screening method, which we have named CRISPRmap. Our method combines a novel in situ CRISPR guide identifying barcode readout approach with concurrent multiplexed immunofluorescence and in situ RNA detection. CRISPRmap barcodes are detected and read out through combinatorial hybridization of DNA oligos, enhancing barcode detection efficiency, while reducing both dependency on third party proprietary sequencing reagents and assay cost. Notably, we conducted a multi-omic base-editing screen in a breast cancer cell line on core DNA damage repair genes involved in the homologous recombination and Fanconi anemia pathways investigating how nucleotide variants in those genes influence DNA damage signaling and cell cycle regulation following treatment with ionizing radiation or DNA damaging agents commonly used for cancer therapy. Approximately a million cells were profiled with our multi-omic approach, providing a comprehensive phenotypic assessment of the functional consequences of the studied variants. CRISPRmap enabled us to pinpoint likely-pathogenic patient-derived mutations that were previously classified as variants of unknown clinical significance. Furthermore, our approach effectively distinguished barcodes of a pooled library in tumor tissue, and we coupled it with cell-type and molecular phenotyping by cyclic immunofluorescence. Multi-omic spatial analysis of how CRISPR-perturbed cells respond to various environmental cues in the tissue context offers the potential to significantly expand our understanding of tissue biology in both health and disease.

## Introduction

Pooled CRISPR screens, where the responses of many individual cells to different genetic perturbations can be measured in parallel, are enabling an increasing variety of high-throughput genetic analyses. All such studies must correlate phenotype with the specific genetic modification, which is generally identified by the readout of a DNA encoded barcode. This was originally achieved for assays of cell viability or expression of a marker used for cell sorting by measuring enrichment or depletion of specific barcodes from the bulk or sorted cell population. To allow screening for molecular phenotypes it is necessary to measure these parameters and associated barcodes in single cells. Coupling single-cell RNA-sequencing to pooled CRISPR screens has vastly expanded our ability to study the transcriptomic response to perturbations^1,2^, but suffers from the necessity to isolate and lyse cells. As such, scRNA-seq approaches are agnostic to spatial organization of inter- and intracellular phenotypes. Imaging techniques have emerged to enhance these screens^3,4^, capturing complex cellular behaviors and dynamic phenotypic changes, including intricate cellular morphology and molecular distribution, without destroying the cells. These advancements make it possible to observe a wide array of spatially resolved cellular phenotypes in genetic screens.

The integration of single-cell multi-omic profiling, which concurrently analyzes proteins and RNA, is crucial for a nuanced understanding of cellular function. While RNA profiling provides data on gene expression, sole reliance upon RNA profiling can be fraught with incomplete or erroneous conclusions due to post-transcriptional and post-translational processes that RNA sequencing is blind to. Combining multimodal profiling with the ability of optical pooled CRISPR screens to perturb at a pathway- or even genome-wide scale, has not yet been widely adopted, despite its potential to expand our understanding of how cellular pathways are regulated in health and disease states like cancer.

Pooled base editor screens have recently emerged as a powerful method for in situ mutational scanning, enabling researchers to directly alter endogenous proteins within live cells and thereby revolutionize the study of proteins in their natural environments^5,6^. These screens can utilize nuclease hybrids of deficient Cas9 with APOBEC1 (BE3) guided by single guide RNAs (sgRNAs) to induce specific point mutations through direct chemical modification, offering a precise means of editing^7^. These precise base changes largely occur within a defined genomic nucleotide window^8^. The precision of editing is advantageous for high-resolution analysis of protein function, and the mapping of sequence-activity relationships.

Leveraging multi-omic optical pooled screens to interrogate key cellular and clinically relevant pathways holds immense potential. Indeed, as next-generation sequencing has become more common in clinical oncology, there has been an increasing number of variants of unknown significance (VUS) in genes linked to cancer predisposition and aggressiveness^9^. In particular, VUS are commonly identified in DNA damage response (DDR) genes, which are critical for genomic stability, DNA damage signaling, DNA-damage related checkpoints, and DNA repair genes^10^. The importance of understanding whether a VUS is functional or a passenger event is underscored by real-world clinical consequences. For example, patients with particular VUS may be a candidate for therapy leveraging impairment of a DDR pathway or whether relatives may be at elevated cancer risk if such a mutation arises in the germline. As such, understanding the normal function of each gene and how key mutations alter homeostasis is critical. Furthermore, due to the essential nature of many DDR genes for cell viability, efforts to genetically deplete DDR genes may not recapitulate clinically-observed variants and function. Hence, efforts to interrogate the function of these proteins with point mutations is necessary to understand their function.

Ideally, the effects of variants, or any genomic perturbation, would be studied in the native context the cell encounters in vivo. Recent technological advancements have allowed for protein epitope-based identification of CRISPR guide expressions within tumor tissues at a single-cell resolution^11,12^. RNA-based barcoding holds the promise to increase the complexity of these libraries, but its application in tissue contexts has not yet been reported.

Building on these innovations, we have developed a novel, sequencing-free barcode readout approach for optical pooled CRISPR screens that is compatible with highly multiplexed antibody and RNA transcript profiling. We have applied this method to a breast cancer cell line to evaluate how 292 nucleotide variants across 27 key DDR genes affect the DNA damage response by visualizing the recruitment of DDR proteins to sites of DNA damage during different cell cycle phases after ionizing radiation exposure. Our work also demonstrates the capability to optically read RNA-encoded barcodes in tissue sections, linked with multiplexed antibody detection. This serves as a stepping stone toward in vivo CRISPR screens that can map the cellular landscape and pathway behaviors at a subcellular level.

## RESULTS

### CRISPRmap enables optical readout of cellular barcodes

Pooled CRISPR screens typically introduce a single perturbation and its corresponding barcode in a cell through lentiviral infection at a low multiplicity of infection (**Figure 1a & b**). Our barcode is expressed as part of an abundant mRNA encoding for a selection marker^3^ (**Figure 1c**). In CRISPRmap, the cellular barcode consists of a unique combination of two adjacent 30 bp hybridization sequences. The first step of barcode detection occurs through hybridization of a pair of ssDNA oligos that are complementary to the adjacent hybridization sequences on the transcript^13^ (**Figure 1e**). In our approach, the primer and padlock oligos each contain a unique pair of 20mer readout sequences. Collectively, the four 20mer sequences form a unique combinatorial readout set. Padlock probe circularization by T4 DNA ligase is dependent on hybridization of splint oligos, which bind to the 20mers on the primer oligo (**Figure 1e**). Subsequently, rolling circle amplification is initiated through the primer oligo. Crucially, valid amplicons rely on AND-logic for the primer, padlock and both splint oligos. The readout set, and thus by extension the cellular barcode, is identified by cyclical hybridization rounds with dye conjugated oligos (readout probes)^14,15^. Distributing the readout set over primer and padlock probe enables us to identify improperly self-ligated padlock oligos (readout set lacks primer readouts), or unallowed primer-padlock pairing (invalid readout set), and exclude them from analysis. Our cyclical hybridization readout approach was designed to minimize dependence on 3^rd^ party sequencing reagents, tissue degradation during cyclic enzymatic steps, and reagent cost of the assay (**Supplementary Table 1**).

**Figure 1.**
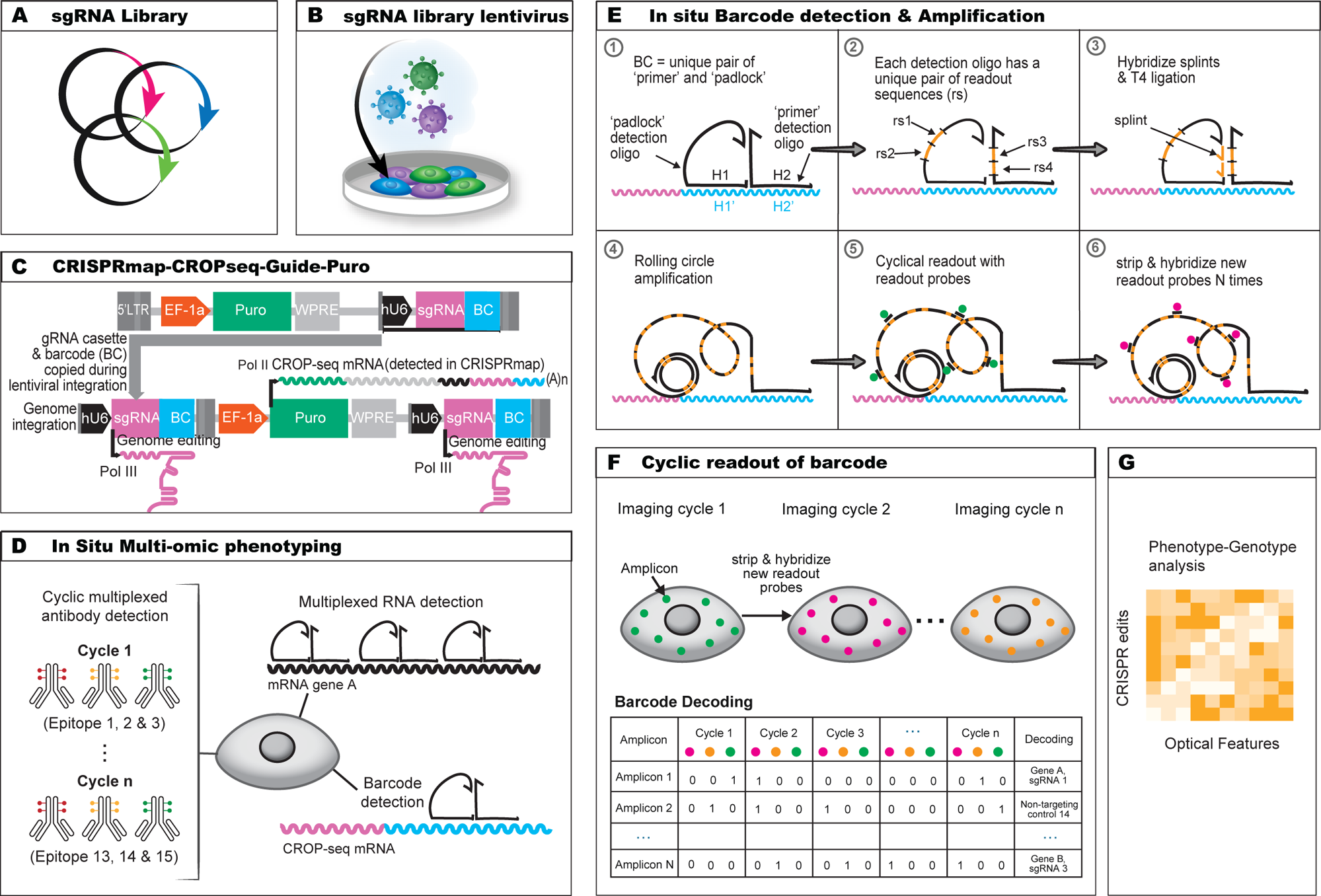
CRISPRmap assay design overview. **(A)** Synthesized single-guide RNA (sgRNA) and barcode (BC) library are cloned onto the modified CROPseq vector for sgRNA and BC expression. **(B)** Plasmids are lentivirally transduced into target cells. **(C)** Design of the CRISP-map-CROPseq-Guide-Puro vector. Human U6 (hU6) promoter (black) drives the sgRNA expression by RNA Pol IIIand a Pol III stop signal is inserted between the sgRNA and the barcode (BC). The hU6-sgRNA-stop cassette and the BC are inserted in the 3’ LTR sequence and will thus be copied during genome integration to the upstream of the EF-1a promoter. The EF-1a promoter drives the expression of the CROPseq mRNA by RNA Pol II, which expresses the Puromycin resistance gene (green), hU6 (black), sgRNA (magenta) and BC (cyan). **(D)** In situ multi-omic phenotyping and CRISPRmap barcode detection. Multi-omic phenotyping interrogates proteomic and transcriptomic states while CRISPRmap barcode readout identifies the sgRNA identity. Cyclic antibody staining (IBEX) is used to detect dozens of epitopes. Pairs of padlock and primer oligos are hybridized to the CROPseq mRNA or endogenous RNAs to detect CRISPRmap barcodes or target RNA transcripts. **(E)** In situ barcode detection and amplification. Padlock and primer oligos hybridize to the BC sequence on the CROP-seq mRNA. Padlock and primer oligos each encode a unique pair of readout sequences (rs). Splints hybridize to the corresponding rs sequences on primer oligos. Padlock oligos and splints are joined by T4 ligation to enable formation of amplicons by rolling circle amplification. Fluorophore-conjugated readout probes hybridize to readout sequences on the amplicons in a cyclic manner for barcode identification. **(F)** Barcode readout and decoding. Images across fluorescence channel and imaging cycles are co-registered into a unified readout stack. Barcode decoding at the amplicon level is achieved through spot detection, assigning a bit code (0 for absence, 1 for presence) in each image to generate a barcode across images. If the barcode aligns with a guide-identifying barcode in the codebook, a guide identity is assigned to the corresponding amplicon. **(G)** Phenotype-genotype analysis. Multi-omic and multiplexed phenotyping provides high dimensional optical features for systematic analysis.

To develop and optimize our approach we transduced a small pilot lentiviral library containing five GFP-targeting CRISPR guides, and five non-targeting CRISPR guides in a HT1080-Cas9 cell line expressing copGFP. We performed lentiviral library preparation (**Methods**) and infected cells at an MOI of 0.1 to ensure most infected cells will be edited by a single guide, and express a single barcode after puromycin selection. To couple our optical phenotype (GFP expression) to the CRISPR edit, we performed CRISPRmap (**Methods, Figure 2a**). For this small pilot experiment, the readout set was imaged in 2 channels over 4 imaging cycles, larger libraries discussed later were imaged in 3 channels over 8 imaging cycles. Images across all barcode readout cycles and channels were co-registered into an image stack and corrected for global translational shifts (i.e. misaligned glass bottom well plate placement) as well as local translational shifts (i.e. cells slightly shifting between imaging rounds). To align the images across all imaging rounds we calculated the transformation matrices for each round using the TV-L1 implementation of optical flow^16^ on binary nuclei masks derived from DAPI stains (**Methods**). In our GFP-targeting CRISPRmap screen barcode decoding is performed at amplicon level (**Methods**), by assigning an 8-bit code for each amplicon across the readout cycles and channels, where signal from each readout sequence yields a positive entry (1), and lack of signal a negative entry (0) (**Figure 2b**). A guide identity (Guide ID) is assigned to an amplicon if the 8-bit code of the amplicon position matches a guide-identifying barcode in the pre-designed library codebook. We found that of all the amplicons that were positive for four readout probes, 98% coded for an allowed barcode included the library design, whereas 2% of amplicons reported an unallowed barcode (**Figure 2c**), despite their relative ratios of 10/25 vs 15/25 possible primer-padlock pairs. From a per-cell analysis, we found that when imaging with a 20x objective, the median number of guide-assigned amplicons per cell was 11 (**Figure 2d**). We restricted further analysis to cells with 3 or more amplicons and for which the most abundant barcode made up more than two thirds of the amplicons of the sum of the two most abundant barcodes under a cell segmentation mask. The latter criterium was put in place to retain cells for which imperfect cell segmentation could cover a few amplicons from neighboring cells, causing false association of guide-assigned amplicons to cell masks. With these quality control metrics in place, we retained 76% of the cells for further analysis (**Figure 2d, Supplementary Table 2**).

**Figure 2.**
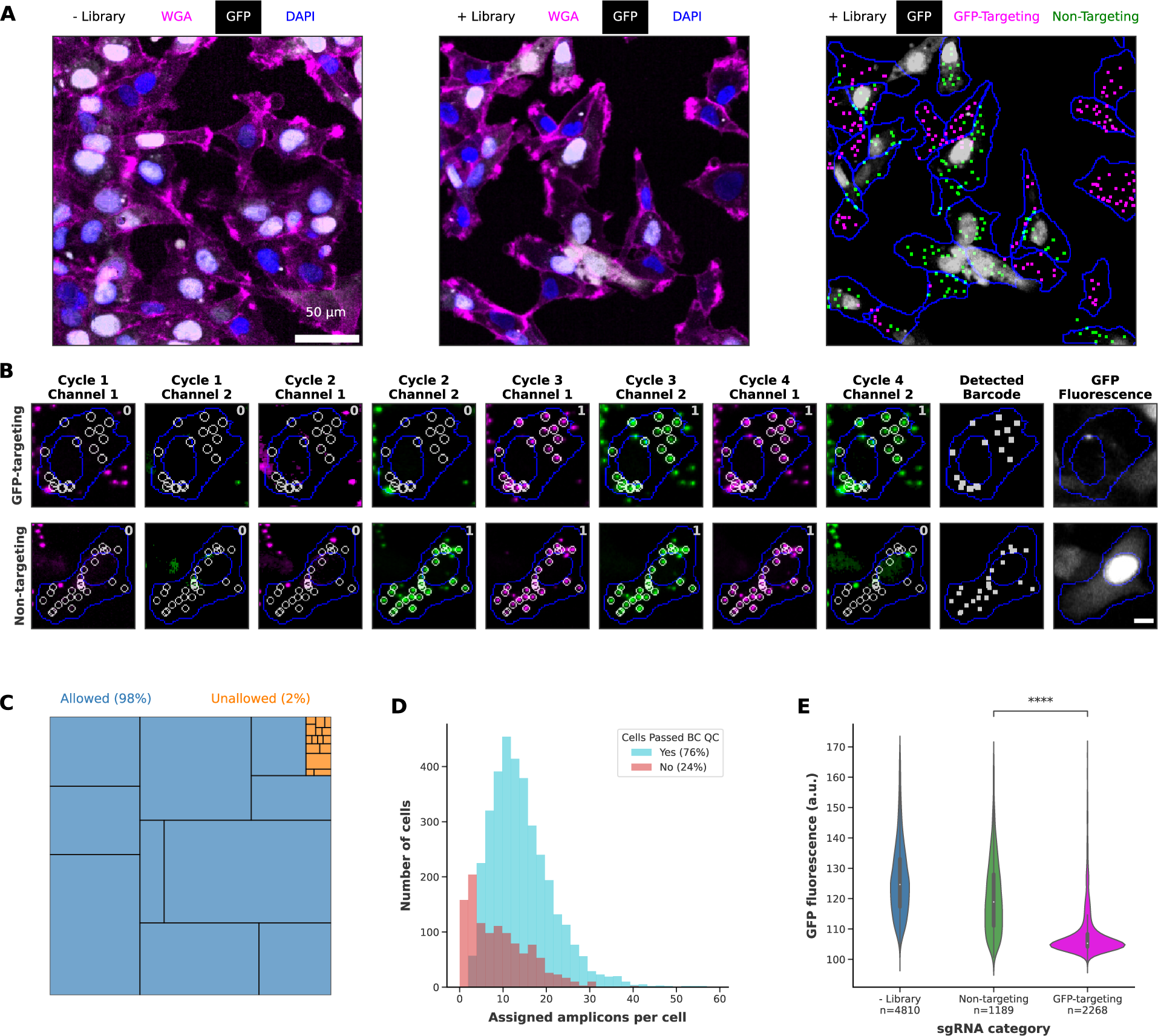
CRISPRmap shows high-fidelity genotype-phenotype mapping. **(A)** Visualization of geno-type-phenotype mapping, showing loss of GFP expression in cells identified with GFP-targeting guides. GFP-expressing HT1080-Cas9 cells without (left) or with (middle and right) GFP-pilot library are shown. In the left and middle panel, WGA stains the cell membrane (magenta) and DAPI stains the nuclei (blue). In the right panel, cell boundaries determined by cell segmentation are outlined in blue, whereas decoded barcodes are shown as false colored spots(magenta for the GFP-targeting guides, green for the non-targeting control guides). Raw GFP signal is displayed by gray-scale on all three panels. Scale bar, 50 µm. **(B)** Visualization of the barcode readout and phenotyping, showing a cell with a GFP-targeting guide (top row, barcode is ‘00001111’) and a cell with a non-targeting control guide (bottom row, barcode is ‘00011110’) over 4 readout cycles and 2 fluorescence channels. Decoded barcodes are displayed as spots false colored in white in the column labeled “Detected Barcode” (second most right), and the positions of the decoded amplicons are projected onto the 8 readout images as white circles (column 1 to 8). Raw readout signals are displayed in magenta for channel 1 and green for channel 2. Raw GFP signal is shown in gray-scale in the column labeled ‘GFP fluorescence” (most right). Cell and nuclear boundaries are outlined in blue in all panels. Scale bar, 20 µm. **(C)** Quantification of all possible primer and padlock combinations detected, showing robust detection of the 10 allowed combinations (blue) encoding the 10 guides in GFP-pilot library (98%) and minimal detection of the 15 unallowed combinations (orange) (2%). **(D)** Distribution of the number of assigned amplicons per cell. Quality check (QC) is performed on each cell based on the number and purity of the most representing guide. 76% of cells passed QC (n = 3457). **(E)** Quantification of genotype-phenotype mapping, showing cells with GFP-targeting guides have significantly reduced GFP fluorescence. Cells with non-targeting control guides show GFP fluorescence close to unperturbed cells. Two-sidedMann-Whitney test, *p < 0.05, **p < 0.01, ***p < 0.001, ****p < 0.0001.

To assess guide representation throughout library preparation, infection and optical readout of the barcodes, we performed NGS sequencing on PCR product of the amplified synthesized DNA oligonucleotide pool, the CRISPRmap-CROPseq plasmid pool, and the genomic DNA from the cells transduced with the library. We further compared the sequencing result to the relative guide abundance of cells with optically identified guide identities, and observed highly correlated guide frequencies between all of these stages (**Supplementary Figure 1a & b**).

Furthermore, we evaluated if we observed the expected optical phenotype for each of the guides in our pilot library and found indeed that cells with GFP-targeting guides have significantly lower GFP fluorescence levels than cells with NTC guides (**Figure 2e**), which in turn have similar GFP fluorescence levels as unperturbed cells. Cells with each of the five GFP-targeting guides showed significantly lower GFP levels than cells with any of the NTC control guides under pairwise comparison (**Supplementary Figure 1c**), thus recapitulating the expected genotype-phenotype relationship. Interestingly, CRISPRmap barcode decoding at the amplicon level led to the detection of some double-transduced cells. These cells express two barcodes, each with a unique spatial pattern (**Supplementary Figure 1d**). Future studies could leverage this feature to study genetic interactions through combinatorial perturbations by infecting pooled libraries at a higher multiplicity of infection.

Finally, apart from the HT1080 cells shown in the GFP pilot study, we have validated CRISPR barcode readout in five additional cell lines to date (**Supplementary Figure 2**), showing the broad compatibility of our assay with different cell types.

### Multi-omic in situ phenotypic profiling of perturbed cell states

To unlock the potential of base editor scanning, we aimed to move beyond cellular fitness as a readout and enable detailed characterization of more complex biological processes. Therefore, we sought to combine base editing approaches with optical, single cell, multiplexed multi-omic approaches, measuring functional responses of dozens of proteins and mRNAs at subcellular resolution. We profiled the proteomic and transcriptomic responses of breast cancer cells to ionizing irradiation, a critical treatment modality for breast cancers, as a function of base-edited variants of 27 core DNA damage repair genes involved in the DNA damage response, homologous recombination and Fanconi anemia pathways. Specifically, a 364 sgRNA library was lentivirally transduced into an MCF7 cell line expressing BE3 (MCF7-BE3 hereafter)^5^. Transduced cells were selected in antibiotic-containing medium for two days and cultured in antibiotics-free medium for another two days, prior to induction of DNA damage by gamma radiation. Six hours after irradiation, cells were chemically fixed and profiled for protein expression, mRNA expression, and CRISPR barcode readout (**Figure 3a, b**). In this dataset, we profiled 226,369 single cells that met all barcode calling quality metrics, resulting in an average coverage of 310 cells per variant in the library in each experimental condition.

**Figure 3.**
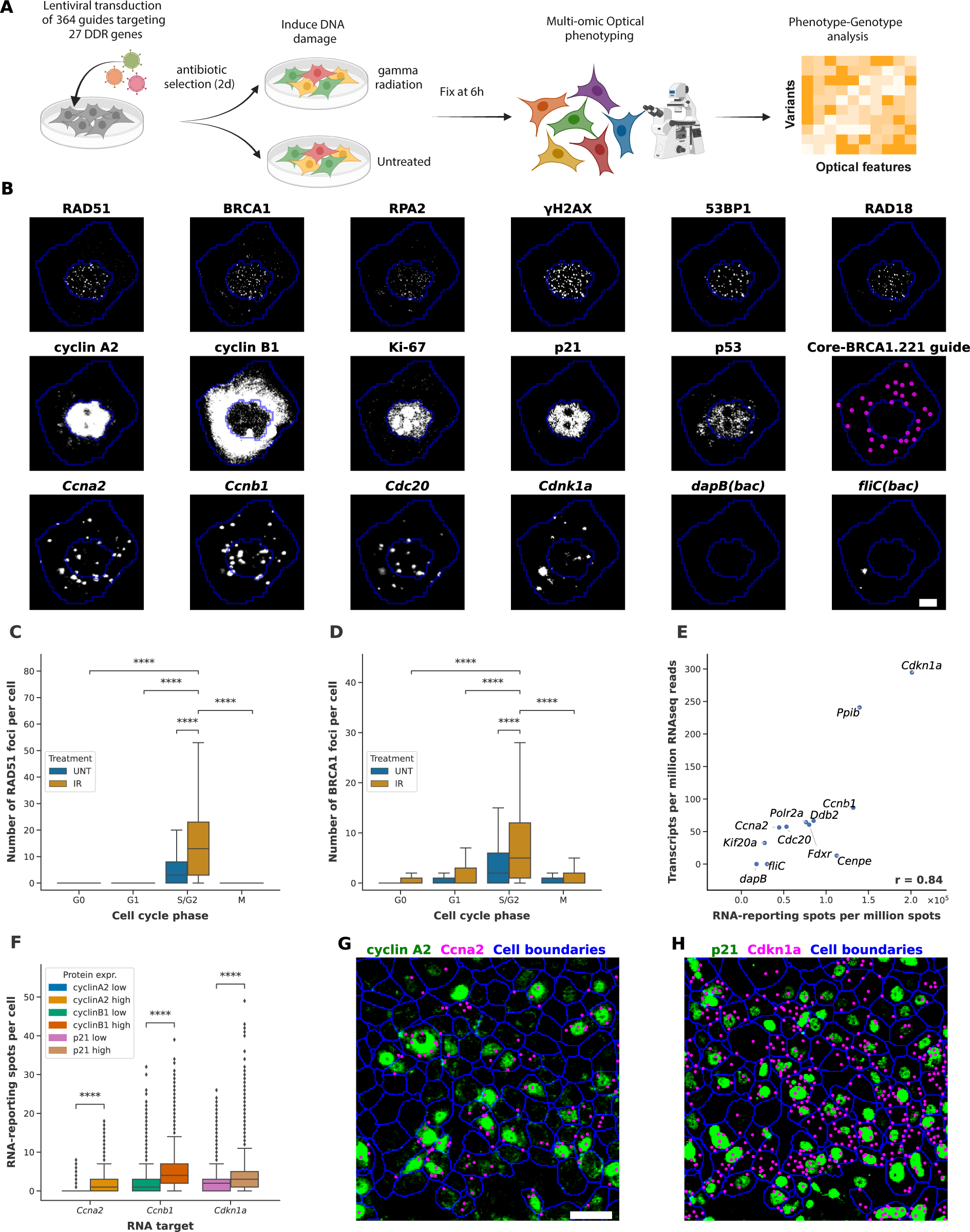
CRISPRmap base-editing screening enables multi-omic phenotyping of cell states. **(A)** Experimental workflow. MCF7-BE3 cells are lentivirally transduced with a 364-guide library targeting 27 core DNA damage response (DDR) genes. After lentiviral infection, cells are selected in puromycin-containing medium for 2 days then cultured 2 days without puromycin. Cells were either gamma irradiated (IR) or kept untreated (UNT), and fixed 6 hours later, prior to multi-omic optical phenotyping and barcode detection. **(B)** The subcellular distribution of 6 DNA damage response protein stains (top row), 5 cell cycle regulator stains (middle row) and barcode detection (middle row, most right), in addition to transcript detection for 6 genes (bottom row) are shown for a single cell. Cell and nuclear segmentation are outlined in blue, whereas raw antibody signal and transcript detection are in gray-scale. Decoded barcodes are shown as false colored (magenta) spots. Scale bar, 10 µm. **(C)** Quantification of the number of RAD51 foci per cell across cell cycle phases in UNT and IR cells, showing significant foci induction by irradiation and enrichment in S/G2 phase. Outliers are omitted in the plot. **(D)** as in C) for BRCA1 foci. **(E)** Correlation between RNA-reporting spots per million spots measured by RNAmap and Transcript Per Million (TPM) reads measured by bulk RNA sequencing. A panel of 12 transcripts were measured. Pearson correlation (r) equals 0.84. **(F)** Quantification of RNA-protein correlation measured by RNAmap and antibody staining for three RNA-protein pairs (Ccna2-cyclin A2, Ccnb1-cyclin B1, and Cdkn1a-p21), showing significant enrichment of RNA-reporting spots in cells with high expression of the encoded protein. **(G)** Visualization of correlated expression of RNA and protein between Ccna2 and cyclin A2. Decoded Ccna2-reporting spots (magenta), and cyclin A2 antibody signal (green) are shown. Scale bar, 50 µm. **(H)** as in G) for Cdkn1a and p21. The same cells as in G) are shown. Two-sided Mann-Whitney test, *p < 0.05, **p < 0.01, ***p < 0.001, ****p < 0.0001

To evaluate how variants alter the cellular response following treatment with ionizing radiation, we applied a recently developed approach IBEX, which employs a cyclical process of antibody staining and chemical bleaching to facilitate high-resolution imaging of dozens of epitopes within a single sample, while preserving its physical integrity^17^. We visualized a panel of key DNA damage response proteins (RAD51, BRCA1, RPA2, γH2AX, 53BP1 and RAD18), cell cycle phase marker proteins (Ki-67, cyclin A2, cyclin B1, and phospho-Histone H3), and apoptosis-related proteins (cleaved PARP1, p21 and p53), and recovered expected subcellular protein localizations (**Figure 3b**) and treatment-specific staining patterns (**Figure 3c, d**). We also quantified the micronuclei formation based on DAPI stain (**Methods**). Accumulation of DNA damage response proteins at damaged genomic loci typically gives rise to punctate immunofluorescence detection patterns (foci) in the nuclei, which we quantify at the single-cell level through automated detection (**Methods, Supplementary Figure 3a**). Cell cycle related proteins and transcription factors on the other hand are evaluated as average fluorescence across the cellular, nuclear and cytosolic mask (the latter we define as cell mask minus nuclear mask). Comparison of cytosolic to nuclear abundance of a protein allows for quantification of its translocation status. To further assess the specificity of the antibody staining, we compared the expression level and subcellular localization of the measured proteins in gamma-irradiated cells to untreated cells, while separating cells into four different cell cycle phases (G0-, G1-, S/G2-, M-phase) based on the cell cycle markers we measured. As expected, we observe significant induction of nuclear foci formation for all the six measured DNA damage response proteins upon gamma irradiation (**Figure 3c, d, Supplementary Figure 3c-f**). In addition, we observe the expected significant enrichment of nuclear foci that function in the homologous recombination pathway (RAD51, BRCA1, RPA2 and RAD18) in the S/G2-phase. Large γH2AX foci are reported to be involved in double-strand break signaling^18^ and 53BP1 foci believed to promote the non-homologous end joining pathway showed a slight enrichment in G1-phase over S/G2 phase (**Supplementary Figure 3e, f**). Finally, the single-cell and highly multiplexed nature of our data enables us to evaluate the correlation among all optical features we measured (**Supplementary Figure 3h**). As expected, we observe a positive correlation between the nuclear foci involved in homologous recombination (RAD51, BRCA1, RPA2 and RAD18) with the average nuclear intensity of the S/G2-phase markers (cyclin A2, cyclin B1) and the proliferation marker (Ki-67).

To simultaneously measure the transcriptomic response of cells, we adapted our CRISPRmap barcode detection approach to detect endogenous mRNA transcripts, and call this approach RNAmap (**Figure 1c**), which differs from CRISPRmap in three key ways. First, the transcript hybridizing regions of the primer and padlock detection oligos target adjacent sequences on the endogenous RNA transcripts. The design of gene-specific detection oligos was refined for specificity, a narrow range of melting temperatures for primer and padlock oligos, minimal off-target binding and secondary structure (**Methods**). Secondly, to promote detection efficiency, we increased the number of primer-padlock oligo pairs to six per RNA transcript. Primer-padlock pairs that share an RNA target also share the same set of readout sequences. Thirdly, to further boost detection efficiency, RNAmap primer detection oligos only encode a single readout sequence, and as a result, only a single splint oligo needs to undergo ligation to form an RCA template. Consequentially, the padlock oligo encodes three readout sequences to enable similar combinatorial readout of transcript identities.

We applied RNAmap to target a panel of 12 genes, selected for their expression in irradiated MCF7-BE3 cells, and to span a range of expression levels. The transcripts profiled include cell cycle-related (*Ccnb1*, *Ccna2*, *Cdkn1a, Cdc20, Kif20a, Cenpe*), housekeeping (*Ppib & Polr2a*), DNA damage response-related (*Ddb2*, *Fdxr*) and bacterial negative control (*dapB*, *fliC*) genes (**Figure 3b**). We validate the specificity of detection by comparing optically identified transcript-reporting spots at the population level to bulk RNA-sequencing reads, and observe a Pearson correlation of 0.84 (**Figure 3e**). Additional support of specificity is provided by the analysis of the correlation between mRNA and protein expression level for the three cell-cycle related genes. Cells for which we observe high abundance (**Supplementary Figure 3c**) of cyclin A2, cyclin B1, and p21 at the protein level, have significantly (p < 0.0001, two-sided Mann–Whitney U test) more corresponding transcripts detected by RNAmap (**Figure 3f-h**).

Of note, the degree of multiplexed phenotypic readout of our approach can be further expanded for both protein and transcriptomic detection. In this study, while we only profiled 12 transcripts, our RNAmap design is extendable to detect the expression of hundreds to thousands of genes, similar to recent hybridization based optical transcriptomic approaches, which have profiled the expression of hundreds to thousands of genes^19–22^. Multiplexed immunofluorescence assays, such as IBEX, have achieved concurrent profiling of ∼60-100 protein targets^17^. Similarly, our approach can easily be scaled up to accommodate larger screening library sizes. The set of 54 primer and 54 padlock oligos used in this study enables up to 2,916 barcodes, but we expect that this can be readily scaled up to enable genome-wide screening.

### CRISPRmap base-editing screen with optical phenotyping recapitulates known DNA damage response pathway mechanisms

Building upon our prior work assessing human nucleotide variants across 86 DDR genes and assessed their effect on cell viability^5^, we selected variants that significantly altered viability in at least one treatment condition in the prior study, we focused the present study on sgRNAs with a single C base in their editing window, to minimize confounding effect of bystander mutations that could obfuscate the phenotypic consequences associated with a guide. Combining the 292 guides targeting DDR genes with 35 guides targeting the AAVS1 safe-harbor site and 37 non-targeting control (NTC) guides that have minimal targets in the human genome, we applied a 364-guide library (referred as DDR364) to the MCF7-BE3 cells for a multi-omic pooled optical base-editing screen. Our library includes 162 missense guides, 50 nonsense guides and 80 splice guides (variants that affect splice-donor or -acceptor sequences). It is expected that nonsense variants and splice variants are more detrimental to protein function than the missense variants, which are associated with a broader range of effects on protein function. Our library includes 64 variants that the ClinVar^23^ database annotates as pathogenic/likely pathogenic (P/LP) variants, and 75 variants with uncertain significance (VUS).

We set out to validate if CRISPRmap could recapitulate known phenotypes as a function of specific base edits. First, similar to our previous fitness study^5^, we found stronger phenotypic changes for guides with higher Rule Set 2 (RS2) on-target efficiency scores, initially created for CRISPR-KO sgRNAs^24^. Specifically, we first evaluated the abundance in RAD51 foci for variants of genes that are essential for RAD51 foci formation (RAD51D, RAD51C, XRCC3, BRCA1, BRCA2) relative to negative control guides. Here, we observe a trend of decreasing RAD51 foci with guides with higher RS2 scores, especially for guides that are from the deleterious (nonsense and splice) categories (**Figure 4a**). Notably, implementing a minimum Rule Set 2 score threshold significantly improved the differentiation between deleterious and negative control guides (two-sided KS-test, p.adj<0.0001, **Figure 4c**), while splice variants below the RS2 threshold show no statistical significance over control guides (two-sided KS-test, p.adj>=0.05, **Figure 4b**). Missense guides targeting these genes generally show much milder impact on the abundance in RAD51 foci. Similarly, phenotypic changes in BRCA1 foci abundance of BRCA1 variants strongly correlated with RS2 scores (Pearson r= −0.75, **Figure 4d**), and cells expressing nonsense or splicing variant guides with RS2 >=0.55 showed significantly fewer BRCA1 foci than cells expressing negative control guides (**Figure 4f**). This distinction was not significant for guides with lower RS2 scores (**Figure 4e**).

**Figure 4.**
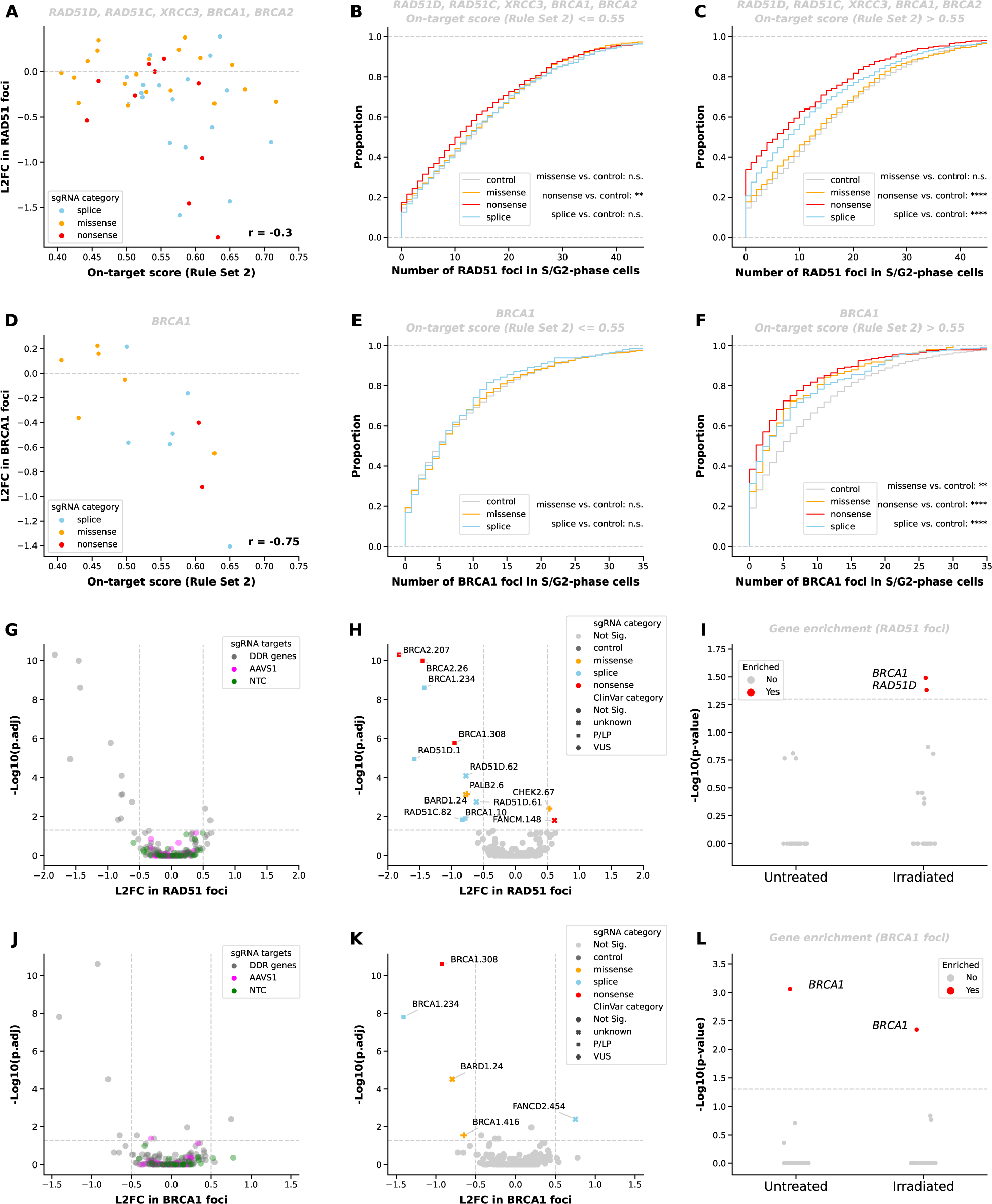
Performance of CRISPRmap base-editing screening on gamma-irradiated MCF7 cells. **(A)** Correlation between log2-fold change (L2FC) in RAD51 foci number and the Rule Set 2 on-target score. All guides targeting RAD51 regulators including RAD51 paralogs (RAD51D, RAD51C, XRCC3), BRCA1, and BRCA2 in the library are shown. Splice and nonsense variants with high Rule Set 2 score shows more significant L2FC. Pearson correlation (r) equals −0.30. **(B)** Quantification of RAD51 foci in irradiated S/G2-phase cells with guides targeting RAD51 regulators that have low Rule Set 2 score. Cells are grouped by sgRNA category. No. or moderate significant separation from the cells with control guides is observed. Two-sided KS test, *p.adj < 0.05, **p.adj < 0.01, ***p.adj < 0.001, ****p.adj < 0.0001. (Outliers are not shown in the CDFs.) **(C)** as in B) for guides with high on-target score, showing significant reduction in RAD51 foci in cells with nonsense and splice guides, compared to the control guides. **(D)** as in A) for L2FC in BRCA1 foci for BRCA1-targeting guides, showing a strong negative correlation with the Rule Set 2 score. Pearson correlation (r) equals −0.75. **(E)** same as B) for BRCA1 foci and guides targeting BRCA1 that have low Rule Set 2 score, showing no significant separation from the control guides. **(F)** same as E) for guides with high Rule Set 2 score, showing significant reduction in BRCA1 foci in cells with missense, nonsense, and splice guides. **(G)** Volcano plot showing no guides targeting AAVS1 and non-targeting control (NTC) guides shows statistically significant changes in RAD51 foci under irradiation. Significance is defined by p.adj < 0.05 and absolute L2FC > 0.5. All guides targeting DDR genes with on-target score >= 0.55, all AAVS1-targeting non-targeting control (NTC) guides are shown. Benjamini-Hochberg corrected two-sided KS test. **(H)** Same as G) showing guides that result in significant changes in RAD51 foci. **(I)** Gene enrichment analysis of guides targeting RAD51D among guides causing significant changes in RAD51 foci in the irradiated condition. Fisher exact test. Enriched genes are defined as p.adj <0.05. **(J)** same as G) for BRCA1 foci. **(K)** same as H) for guides that result in significant changes in BRCA1 foci **(L)** same as I) for BRCA1 foci.

Secondly, we considered variants to significantly alter nuclear foci formation if there was an absolute log2-fold change (L2FC) > 0.5 in the mean number of foci per cell when compared to the population of negative control guides, and we observed a Benjamini-Hochberg corrected two-sided KS-test p-value (p.adj) < 0.05. Based on these criteria, we observe that none of the negative control guides (AAVS1-targeting or NTC guides) significantly alter the number of RAD51 or BRCA1 foci (**Figure 4g & j**). Most of the hits significantly reducing the number of RAD51 foci are nonsense or splice variants, annotated in ClinVar as pathogenic or likely pathogenic (P/LP), but we also identify two missense variants that are annotated as variants with uncertain significance (VUS) or not documented (unknown) in the ClinVar database (**Figure 4h**). Furthermore, guides that target the BRCA1 and RAD51D genes are significantly enriched in the guides that lead to significant changes in RAD51 foci in the irradiated cells (**Figure 4i**), whereas variants that are identified to significantly change the BRCA1 foci (**Figure 4k**) are enriched for BRCA1 variants (**Figure 4l**), as expected. A BRCA1 missense VUS, and an unknown BARD1 missense variant were identified to significantly reduce BRCA1 foci in irradiated cells. The hits identified based on L2FC and p.adj thresholds were further confirmed by the measurement of bootstrapped Wasserstein distance (**Methods**) of each guide from the cells with control guides for RAD51 foci (**Supplementary Figure 4a**) and BRCA1 foci **(Supplementary Figure 4b**).

Apart from nuclear foci, we also analyzed protein stains such as cyclin A2, Ki-67 and p21, for which cells can be categorized into high or low protein expression categories based on the average nuclear fluorescence of the protein stain. We performed beta binomial testing (**Methods**) and observe that three BRCA1 splice variants significantly upregulated the proportion of cells with high p21 expression in the untreated cells **(Supplementary Figure 4c, left)** whereas two ATM splice variants reduced the proportion of cells with high p21 expression in irradiated cells **(Supplementary Figure 4c, right**).

To further verify perturbation-specific phenotypic changes observed in the pooled screens, we selected 2 AAVS1-targeting control guides and 7 guides targeting BRCA1, BRCA2, BARD1, and PALB2 from different sgRNA and ClinVar categories and with on-target score (RS2) > 0.5: (**Supplementary Table 8**). BRCA1.234 (splice, P/LP) and BRCA2.207 (nonsense, P/LP) were identified as hits for RAD51 foci changes in the pooled screen, whereas BRCA1.234 (splice, P/LP) and BRCA1.416 (missense, VUS) as hits for BRCA1 foci changes. We transduced MCF7-BE3 cells with each guide individually, and measured the L2FC in RAD51 and BRCA1 foci compared to all control guides in the gamma-irradiated cells. High correlations of L2FC are observed between guides transduced in the pooled library and transduced individually (Pearson r= 0.90, **Supplementary Figure 4d**; Pearson r= 0.95, **Supplementary Figure 4e**), with hits identified in the pooled screening also showing the most significant changes when transduced individually.

We also applied the same DDR364 library on MCF7-BE3 treated with DNA damaging agents (Camptothecin, CPT; Olaparib, OLAP; Cisplatin, CISP; Etoposide, ETOP) to study the treatment-specific responses of the gene variants (**Supplementary Figure 4f**) with a primary focus on the formation of RAD51 and large γH2AX foci. Similar to the ionizing irradiation screen, we observed that guides targeting the RAD51-relevant genes show more significant loss in RAD51 foci with a higher RS2 score, especially in cells treated with CISP and OLAP (**Supplementary Figure 4g & h**). A RS2 score threshold of 0.5 distinguishes deleterious guides that show mild and strong phenotypes. Again, we observe most of the hits reducing the RAD51 foci in CISP- and OLAP-treated cells are splice or nonsense variants, many of which being P/LP (**Supplementary Figure 4i & j**), and a significant enrichment of guides targeting RAD51 paralog genes, BRCA1, or BRCA2 can be seen (**Supplementary Figure 4k**). A drastic difference in the hits can be seen when comparing the change in large γH2AX foci in ETOP-treated (**Supplementary Figure 4l**) and OLAP-treated (**Supplementary Figure 4m**) cells. In ETOP-treated cells, we observe most splice variants of ATM significantly reduced the number of large γH2AX foci, whereas many deleterious variants coming from a mixed background of RAD51-related genes, ATR, FA pathway genes, increased large γH2AX foci under other treatments. The enrichment test shows a significant enrichment of FANCA- and FANCI-targeting guides in OLAP-treated cells and ATM-targeting guides in ETOP-treated cells among hits of large γH2AX foci (**Supplementary Figure 4n**).

Collectively, these results underscore that CRISPRmap effectively couples barcodes to their corresponding guides, and that the expected phenotypic changes are detected by profiling a few hundred cells per variant, despite imperfect efficiencies associated with current base editors. Moreover, CRISPRmap enables the dissection of therapeutic treatment-specific responses of known pathogenic variants and the identification of novel variants with significant phenotypic effects in the DDR pathways.

### CRISPRmap base-editing screens correlate variants of uncertain significance with known pathogenic variants based on optical phenotypes

Interpreting the functional implications of somatic mutations in cancer, primarily characterized by single nucleotide changes that often lead to missense VUS variants remains a challenging endeavor. This challenge poses a barrier to effective diagnosis, patient stratification, and the management of drug-resistant diseases. Utilizing experimental approaches is crucial in assessing the functional impact of VUS. This is essential for establishing a link between VUS and disease-related phenotypes, particularly due to the limited availability of clinical datasets, and the infrequent occurrence of certain variants in patient cohorts.

We set out to chart variant effects on the DDR pathways by combining the two CRISPRmap base-editing screens that treated cells with ionizing radiation or 4 DNA damaging agents commonly used for cancer therapy. In total, we profiled 948,604 cells that passed barcode QC metrics, averaging 372 cells per guide in each treatment condition. We subsequently clustered variants from functionally related genes (**Figure 5a**) based on the optical features measured in the ionizing irradiation (IR) and DNA damaging agents (CPT, OLAP, CISP, ETOP)-treated cells.

**Figure 5.**
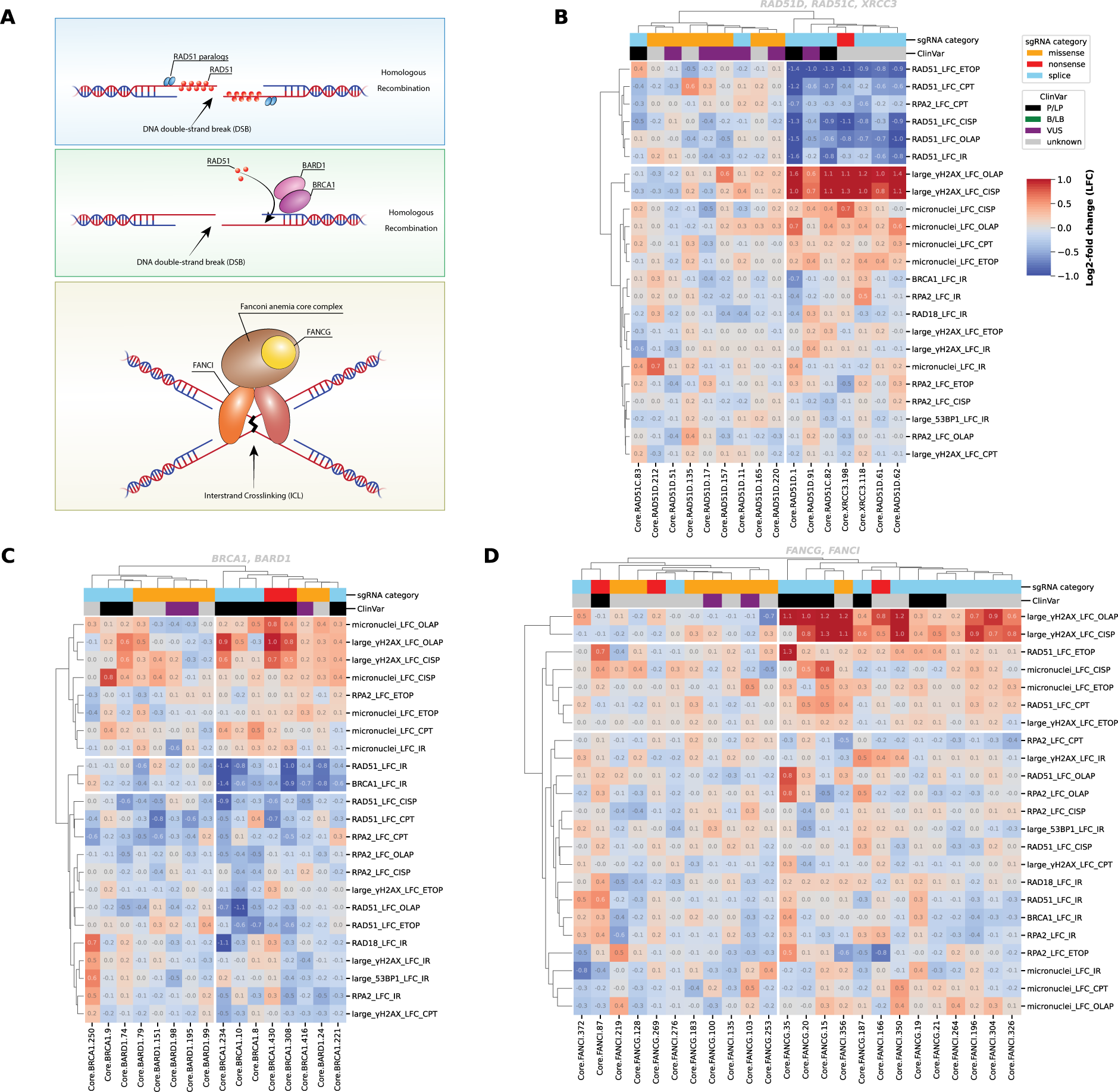
Variant analysis on functionally relevant genes identifies variant clusters with treatment-specific optical signatures. **(A)** Crucial effectors in DNA damage repair. RAD51 paralogs including RAD51D, RAD51C and XRCC3 are required for the formation of RAD51 foci at DNA double-strand breaks (DSBs). The BRCA1-BARD1 complex recruits RAD51 to DSB sites. FANCG and FANCI are involved in DNA interstrand crosslinking (ICL) repair. **(B)** Clustering of guides targeting RAD51 paralog genes (RAD51C, RAD51D, XRCC3), showing a cluster with reduced RAD51 foci in all four DNA-damaging agents-treated and irradiated (IR) cells and increased large γH2AX foci in OLAP- and CISP-treated cells. The mutations of this cluster are mostly splice and nonsense variants. The leftmost column cluster features milder phenotypes mainly associated with missense VUS variants. Log2-fold change (LFC) in each optical phenotype in corresponding treatment conditions were shown as rows in the heatmap. Cells in all cell cycle phases are included, untreated cells are not included. All guides with Rule Set 2 on-target score >= 0.5 were included in the clustering and shown in the heatmap. Columns were cut at a depth of 2 and rows were cut at a depth of 3 based on the dendrogram. Values of L2FC are displayed on the heatmap. Color scale is −1 to 1. **(C)** same as B) for guides targeting BRCA1 and BARD1, showing a cluster with reduced RAD51 and BRCA1 foci in irradiated cells and increased large γH2AX foci and micronuclei in OLAP- and CISP-treated cells composed mainly of pathogenic splice and nonsense variants, and another cluster with mild phenotypes composed mainly of missense variants. **(D)** same as B) for guides targeting FANCI and FANCG, showing a cluster with increased large γH2AX foci and micronuclei in OLAP- and CISP-treated cells. The mutations of this cluster are mostly splicing variants. The leftmost column cluster features milder phenotypes mainly associated with missense variants.

When we clustered the variants from the RAD51 paralog genes (RAD51C, RAD51D and XRCC3), we observed a cluster composed of most splice and nonsense variants (**Figure 5b**, right cluster). This cluster showed a reduction in RAD51 foci across all treatment conditions and a strong upregulation of γH2AX foci for OLAP and CISP treatments, whereas most missense variants form a separate cluster with more mild phenotypic changes (**Figure 5b**, left cluster).

Variants of BRCA1 and its heterodimeric binding partner BARD1 can also be categorized into two clusters. The right cluster contains most splice and nonsense P/LP variants and shows an expected reduction in RAD51 and BRCA1 foci upon irradiation, as well as an increase in large γH2AX foci and micronuclei mostly in OLAP and CISP-treated cells (**Figure 5c**). Variants in the left cluster are mostly missense mutations with milder phenotypes. Notably, a missense VUS variant of BRCA1 (BRCA1.416), that renders the H1283Y amino acid change on the BRCA1 protein, clusters with pathogenic variants of BRCA1. Despite being classified as VUS, a recent study^25^ classified it to be likely pathogenic based on a BE3 base-editing screen with fitness as a readout, and they confirmed the loss in cell viability for H1283Y with CRISPR-mediated homology-directed repair. To assess if the missense BRCA1 VUS variant had a phenotype similar to a nonsense mutation due to a change in protein stability, we performed immunoblotting on cells transduced with individual guides, and found the missense variant to have full length protein at similar abundances as the AAVS1 control variant (AAVS1.86), whereas the splice variant (BRCA1.234) shows a reduction in full-length protein similar to the siRNA-induced BRCA (siBRCA1) knockdown control (**Supplementary Figure 5a**). We also performed immunoblotting on two BRCA2 variants to further investigate the protein stability of different types of variants and we observe a loss of full-length BRCA2 protein in the nonsense variant (BRCA2.207) similar to the siRNA-induced BRCA2 (siBRCA2) knockdown control, whereas the missense VUS variant (BRCA2.438) has similar full-length protein to the AAVS1 control variant (AAVS1.86) (**Supplementary Figure 5b**). Immunoblotting on phospho-KAP1 (pKAP1) indicates the induction of DNA damage upon gamma irradiation and we observe no correlation between the changes in protein stability and the DNA damage (**Supplementary Figure 5a, b**).

Evaluating variants of Fanconi anemia complementation group (FANC) members FANCI and FANCG reveals a cluster (**Figure 5d**, right) that predominantly consists of splice and nonsense variants, with the exception of a single missense FANCI variant (FANCI.356) that increases γH2AX foci for OLAP and CISP treatments far more strongly than other FANC missense variants. Variant clusters with notable signatures can also be found in the clustering outcomes of other functionally related genes, such as BRCA2 and PALB2 (**Supplementary Figure 6a**) and ATM (**Supplementary Figure 6b**). Clustering of all FANC gene variants also identifies clusters with one cluster containing most splice and nonsense variants, showing a similar optical feature signature as in the FANCI-FANCG clustering result (**Supplementary Figure 6c**, left). Besides the missense variants of FANCI, two additional missense variants of FANCM (FANCM.14) and FANCL (FANCL.24) were observed with a similar optical feature signature.

As a whole, these data support that CRISPRmap not only empowers the analysis of missense VUS variants by their correlation with known pathogenic splice or nonsense variants on functionally related genes but also characterizes the drugs-specific responses of these gene variants for multiple DDR pathway regulators. The high-multiplexed nature of CRISPRmap phenotyping further identifies unique optical signatures of variant clusters that sheds light on the molecular mechanisms of known pathogenic variants as well as their correlated VUS variants, making CRISPRmap a potential tool to advance patient-specific precision medicine strategies.

### CRISPRmap enables barcode detection and multiplexed immunofluorescence in tumors

To evaluate if we could read CRISPRmap barcodes in a tissue context, we performed a pilot study and transduced Cas9 negative OE19 (human oesophageal carcinoma) cells with the aforementioned DDR lentiviral library. Following antibiotic selection and expansion, 5 million cells were suspended in a 1:1 mixture of matrigel and PBS and inoculated into the flanks of nude mice. Tumors were harvested after 17 days and flash frozen (**Figure 6a**). The CRISPRmap protocol was performed on sectioned tissue, with minor modifications (**Methods**) to yield ample barcode detection throughout the section (**Figure 6c**) in recognizable epithelial growth patterns, suggesting that single cells yielded local clonal outgrowth. Subsequent multiplexed immunofluorescence profiling enabled cell and nucleus segmentation (**Figure 6c**) based on E-cadherin **(Figure 6h)** and DAPI stains respectively. Despite imperfect cell segmentation of the morphologically diverse tumor cells, and a lack of antibiotic selective pressure during 17 days of tumor growth, the evaluation of barcode signal for segmented cells revealed that 56% of segmented cells pass our barcode QC metrics, and the median number of barcodes detected per cell is 14 (**Figure 6B, Supplementary Table 3**). These metrics are on par with in vitro data. Further antibody staining cycles allowed for the visualization of angiogenesis (CD31, endothelial marker, **Figure 6c, f**), extracellular matrix formation (Tenascin C, **Figure 6c, f**) around tumor domains, as well as a layer of cells expressing vimentin (**Figure 6g)** and N-cadherin (**Figure 6h**), and transcription factor nuclear translocation in the transplanted cells (p21, **Figure 6g**).

**Figure 6.**
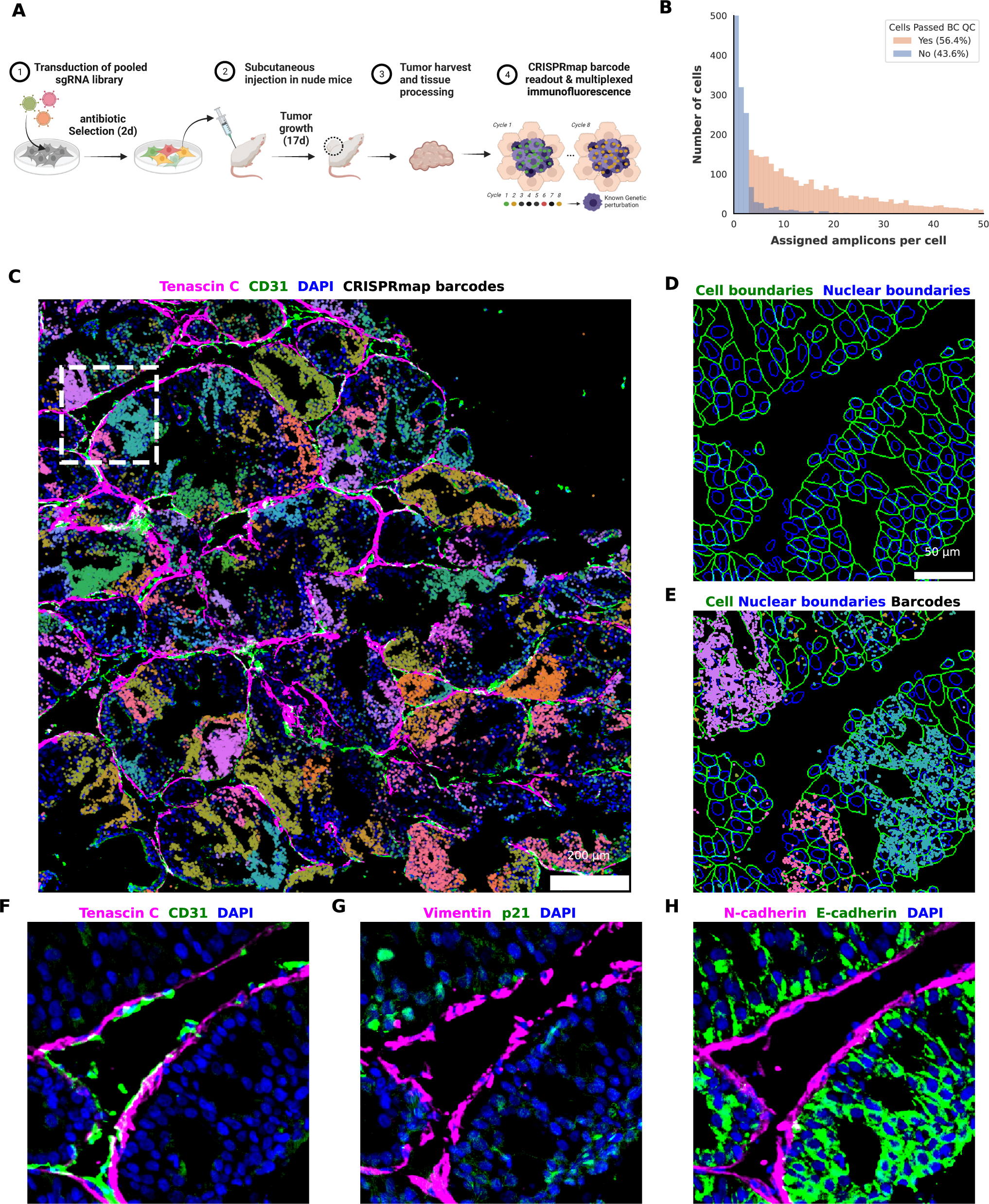
Subcellular resolution CRISPRmap barcode readout and multiplexed phenotyping in vivo. **(A)** Experimental workflow. Cas9 negative OE19 cells were transduced with the 364 guide DDR library and selected with puromycin for 2 days, prior to inoculation in the flank of nude mice. Tumors were harvested after 17 days of growth and processed for CRISPRmap and immunofluorescence imaging. **(B)** Quantification of cells passing the barcode Quality Check (QC) criteria (Methods) for decoded barcodes, showing on par barcode detection with in vitro data. **(C)** Visualization of in vivo iterative immunofluorescence and barcode detection, showing the guide distribution landscape in a tumor section. Extracellular matrix marker Tenascin C (magenta) and mouse endothelial cell marker CD31 (green) line the tumor domains. Decoded barcodes are shown as spots, false colored according to their guide identity. The region highlighted by a white dashed square is zoomed in on D) -H). Scale bar, 200 µm. **(D)** In vivo cell segmentation. Cell (green) and nuclear (blue) boundaries detected by segmentation of E-cadherin and DAPI, respectively. Scale bar, 50 µm. **(E)** Subcellular resolution of barcode readout. Decoded barcodes are shown as spots, false colored according to their guide identity. Cell and nuclear boundaries are shown as in D). **(F)** Iterative immunofluorescence distinguishes cell types and cellular states in vivo. Protein stains of Tenascin C (magenta) and mouse CD31 (green), and DAPI (blue) are shown. Antibodies are predicted to recognize epitopes from both human and mouse origins unless otherwise specified. **(G)** as in F) for Vimentin (magenta) and human p21 (green). **(H)** as in F) for N-cadherin (magenta) and E-cadherin (green).

Iterative immunofluorescence and optically resolved transcriptomics approaches have been established for comprehensive profiling of intracellular, cellular, extracellular and signaling mechanisms in the tumor microenvironment. Coupling spatial genomics to CRISPRmap thus enables in vivo CRISPR screens at subcellular resolutions to systematically interrogate how genomic alterations and protein function modify cellular behavior in a tissue context and/or effects upon the cellular microenvironment. Moreover, CRISPRmap is a CRISPR enzyme agnostic barcode readout, and thus adaptable to the wider CRISPR toolkit, including base editing, gene activation and epigenetic modifications, enhancing the potential to uncover the influence of genes and their regulatory mechanisms on cellular organization and microenvironment.

## DISCUSSION

Our study introduces CRISPRmap, a novel approach in the landscape of functional genomics. By employing a sequencing-free in situ CRISPR barcode readout coupled with cyclic immunofluorescence and in situ RNA detection, CRISPRmap expands upon traditional boundaries of optical pooled genetic screening. This study enriches our understanding of gene functions by enabling systematic examination of spatial phenotypes within perturbed cells—attributes like morphology and subcellular localization of proteins that are lost in sequencing-based methodologies.

Our novel approach reduces the costs associated with complex assays by minimizing reliance on proprietary sequencing reagents. Moreover, CRISPRmap is flexible in the choice of readout dyes so they can be matched to existing microscopy setups available to researchers, encouraging broad adoption in the community.

We applied CRISPRmap to investigate the functional consequences of nucleotide variants in genes critical for the DNA damage response. Our multi-omic profiling of approximately 1 million cells enables a nuanced interrogation of how these variants influence cellular response to DNA damage by expanding the phenotypic profiling from fitness or a single fluorescent reporter to measuring dozens of DDR genes at the proteomic and transcriptomic level with subcellular spatial resolution. This enhanced view of the DDR response empowered us to identify missense variants of uncertain clinical significance whose DDR response resembles known pathogenic nonsense or splicing variants more closely than most VUS or unknown missense variants. As such, our approach can provide a novel framework for annotation of human variants in a treatment specific manner, and can help prioritize therapeutic strategies.

Recently, an elegant use of triplet combinations of linear epitopes enabled antibody-based identification of CRISPR guide expression of dozens of different guide expressing cancer cell populations within a tumor tissue at single-cell resolution and tissue scale^11,12^. RNA based barcoding offers an opportunity to increase library complexity to genome scale^26^, but to date has not been established in a tissue context. As reported in this study, CRISPRmap can detect ample barcode signal in a tissue context with subcellular resolution, which enables the interrogation of cells with their neighbors, and their surrounding extracellular milieu as a function of precise genome editing. Another opportunity for RNA based barcoding is to start exploring the effects of combinatorial gene perturbations, enabled by the spatial resolution of the barcode which offers the capability to detect more than one barcode expressed per cell. This capability could be helpful for screens to prioritize combinatorial targets, and inform therapy modalities that go beyond single-therapy strategies.

However, our study is not without limitations. RNA-based barcoding approaches are reliant on the stability of RNA molecules, which can be challenging in a tissue context due to the presence of RNAse enzymes. A possible resolution for this reliance is to transcribe barcode carrying RNA in vitro by T7 polymerases post hoc, as was recently reported for in vitro cells^27^. This approach enables deep phenotypic profiling of tissue prior to in vitro generation of barcode carrying RNA molecules that can subsequently be detected by CRISPRmap. Moreover, although our RNAmap profiling approach has demonstrated good specificity, the detection efficiency is lower than traditional smFISH approaches. RNAmap allows for strong signal amplification, which enables high throughput imaging (20x objective and short exposure times) thus enabling visualizing millions of cells needed for large scale screening. For any study at hand, it would be of interest to evaluate the balance of deeper transcriptomic profiling enabled by FISH based approaches and throughput and scale of the screen. We expect CRISPRmap to be fully compatible with FISH based transcriptomic profiling. These constraints highlight the need for continual refinement of optical screening techniques and computational analysis methods.

Future directions could focus on expanding the versatility of CRISPRmap to include a broader range of CRISPR modalities, cell types and tissue environments. Studies can also delve deeper into the impact of genetic perturbations on tissue architecture and the interplay between cells in complex microenvironments. We envision that CRISPRmap will pave the way for high-throughput investigations of gene function in diverse biological contexts, from developmental biology to the study of disease pathogenesis.

In conclusion, CRISPRmap marks a significant stride in the field of functional genomics, offering a new lens through which we can examine the intricate tapestry of gene function within and across cells. Our findings herald a shift towards more spatially and temporally resolved studies of gene function, especially in tissue environments, potentially illuminating new paths in precision medicine and the quest to understand the underpinnings of complex biological systems.

## Author Contributions

J.T.G and J.G conceived the GFP KO pilot study. J.T.G., A.C and J.G conceived the base-editing study. J.T.G, E.M.C. and J.G. conceived the tissue profiling study.

J.T.G conceived CRISPRmap and RNAmap, J.G. implemented CRISPRmap.

J.G., Y.S., G.L., and A.T. performed experiments under the supervision of A.C. and J.T.G.

S.H., K.J. and L.P. performed mouse work under supervision of E.M.C.

A.I, B.W., J.G. and N.K. developed the image analysis pipeline under supervision of J.T.G.

S.N. developed the FISHprobe pipeline under supervision of D.A.L. and J.T.G.

J.G., A.I., Y.J., J.Y.Z. and J.H. performed analysis under supervision of E.A., A.C, and J.T.G

J.G., B.W., N.K., Y.J. and J.H. generated figures under supervision of E.A., A.C and J.T.G.

J.G. and J.T.G wrote the manuscript with input from all authors.

## Acknowledgements

We would like to thank Nicoletta Barolini for figure design. We acknowledge support by NIH NCI grants 1DP2CA281605 to J.T.G, R21HG012639 to E.A. and NIH NHGRI grant R01HG012875 to E.A., and NIH grants R01CA197774 and R01CA227450, as well as a Basser Research Grant to A.C.

## Inclusion and diversity

We support inclusive, diverse, and equitable conduct of research.

## Materials and methods

### Cell lines and cell culture

HEK293FT cells (Thermo Fisher Scientific R70007, RRID:CVCL_6911) were cultured in DMEM (Gibco 11965092) supplemented with heat-inactivated 10% fetal bovine serum (FBS) (ATCC 30-2020) and 100 U/ml penicillin-streptomycin (Thermo Fisher Scientific 15140163). To generate constitutive BE3-expressing cells, MCF7 (ATCC HTB-22, RRID:CVCL_0031) cells were transduced with previously generated plasmid BE3-FNLS-P2A-BlastR^28^. MCF7-BE3 cells were cultured in the same medium supplemented with 2 μg/ml Blasticidin (Thermo Fisher Scientific A1113903), HT1080/Cas9 AAVS1 (Genecopoeia SL512, RRID:CVCL_C9F8) cells were cultured in the same medium supplemented with 200 μg/ml HygromycinGold B (Invivogen ant-hg-2). To generate cells with constitutive BFP expression, OE19 (Sigma-Aldrich 96071721, RRID:CVCL_1622) cells were transduced with the plasmid pLV-EF1a-TagBFP2. OE19-BFP cells were cultured in RPMI-1640 Medium (ATCC 30-2001) supplemented with 10% heat– inactivated FBS, 100 U/ml penicillin-streptomycin, 1x Glutamax supplement (Thermo Fisher Scientific 35050079) and 2 μg/ml Blasticidin.

### Library design and cloning

GFP-targeting guides in the GFP-targeting CRISPRmap knockout screen library (referred as GFP-pilot) were designed with CRISPick^29^ by selecting the top 5 recommended candidates in the CRISPRko mode that target the copGFP sequence. The library also contains five non-targeting control (NTC) guides that lack targets in the human genome. Each guide was combined with a universal scaffold sequence and a pair of guide-specific CRISPRmap barcode sequences. Universal 5’ and 3’ homology sequences were then added to facilitate NEB HIFI assembly into the expression vector. Full-length GFP-pilot library sequences are shown in **Supplementary Table 4**. Base-editing guides in the DNA damage response screen library (referred as DDR364) were selected from the base-editing screens as previously described^5^. The library contains 162 missense guides and 50 nonsense guides with a single C base in the editing window (4^th^ to 8^th^ base in the guide targeting sequence), 80 splice-donor or splice-acceptor (referred as splice) guides, 35 guides targeting the AAVS1 safe-harbor site, and 37 non-targeting control guides that have minimal targets in the human genome. All the selected missense, nonsense and splice guides have FDR<0.05 in at least one treatment in the previous screen^5^. Similarly, each guide was combined with the scaffold, CRISPRmap barcode, and homology sequences. Sequences are shown in **Supplementary Table 5**. Both libraries were ordered as synthesized oligo pools (Integrated DNA Technologies) and PCR-amplified with Q5 DNA polymerase (New England Biolabs M0492) using an optimized two-round amplification strategy to minimize barcode-sgRNA recombination, as previously described^30^. Briefly, oligo pools were diluted in ultrapure water (Thermo Fisher Scientific 10-977-023) and 1 pg of total DNA was added to each 50 μl Q5 reaction mix to perform the first-round amplification of 15 PCR cycles, 0.5 μl PCR product from each 50 μl first-round reaction was then added to each 50 μl Q5 reaction mix for the second-round amplification of 10 cycles. Final PCR product was purified with DNA clean & concentrator (Zymo Research D4013). The primer pairs CRISPRmap-F and CRISPRmap-R in **Supplementary Table 6** were used in both rounds. Amplified oligo pools were cloned into a modified CROPseq-puro-v2 (Addgene #127458) vector that removed the original scaffold sequence (referred as CRISPRmap-CROPseq) using NEBuilder HiFi DNA Assembly (New England Biolabs E2621). Next, we electroporated into MegaX DH10B electrocompetent Cells (Thermo Fisher Scientific C640003). An average number of 300 colonies per guide was maintained to better preserve the relative abundance of guides in the library. Bacterial colonies were scraped from the agar plates and pooled together for plasmid extraction (Zymo Research D4212).

### Lentivirus production

293FT cells were seeded into 6-well tissue culture-treated plates at a density of 100,000 cells/cm^2^. After 24 hours, cells were transfected with pMD2.G (Addgene #12259), psPAX2 (Addgene #12260), and CRISPRmap library plasmid (2:3:4 ratio by mass) using Lipofectamine 3000 (Thermo Fisher Scientific L3000001) in lentiviral packaging medium (Opti-MEM (Thermo Fisher Scientific 31-985-070) supplemented with 5% FBS). Media was exchanged after 6 hours and supplemented with 1.5 mM caffeine (Sigma-Aldrich C0750) to increase viral titer. Viral supernatant was harvested at 24 hours and 48 hours after transfection, filtered through 0.45 μm cellulose acetate filters (Corning 431220) and stored in −80°C freezer in aliquots.

### Fluorescence microscopy

All imaging datasets were acquired using a confocal spinning disk microscope (Andor Dragonfly) coupled to a Nikon Ti-2 inverted epifluorescence microscope with automated stage control, Nikon Perfect Focus System, and a Zyla PLUS 4.2Megapixel USB3 camera. Illumination was done with 100mW 405nm, 50mW 488nm, 50 mW 561nm, 140mW 640nm and 100mW 785nm solid state lasers. All hardware was controlled using Andor Fusion software. All images were taken with a NA 0.80 CFI60 Plan Apochromat Lambda D 20x objective (Nikon MRD70270), except for the foci features taken in the DDR364 base-editing screens. Foci features (RAD51, BRCA1, RPA2, γH2AX, 53BP1 and RAD18) and EPCAM in the DDR364-irradaition screen and foci features (RAD51, γH2AX, and RPA2) and EPCAM in the DDR364-chemo screen were taken with a NA 1.4 CFI60 Plan Apochromat Lambda 60x oil immersion objective (Nikon MRD01605). Lasers, laser powers, exposure times, objectives and experiment-specific acquisition parameters are summarized in **Supplementary Table 4 & 5**. Images were acquired with 4 z-slices at 1.5 μm intervals for the cultured cells and with 6 z-slices at 1.5 μm intervals for the tissue sections unless otherwise specified.

### Oligonucleotide fluorophore conjugation

In each 10 μl reaction, 2 μl of 0.5 mM 5’ amine-modified DNA probes (Integrated DNA technologies) is mixed with 1 μl of 10 mM ATTO488-NHS ester (ATTO-TEC AD 488-31), ATTO 643-NHS ester (ATTO-TEC AD 643-31) or CF568 succinimidyl ester (Sigma-Aldrich SCJ4600027) in 1x BBS (Thermo Fisher Scientific 28384) pH 8.5, and incubate at room temperature for 4 hours. Fluorophore-conjugated DNA probes were purified with Oligo Clean & Concentrator (Zymo Research D4060), and diluted to 1μM in ultrapure water, aliquoted and stored at −20C. Oligonucleotide sequences and fluorophores used in the GFP-targeting screen were listed in **Supplementary Table 4**, the base-editing screens and in vivo CRISPRmap barcode readout were listed in **Supplementary Table 5**.

### Antibody fluorophore conjugation

In each conjugation reaction, 5 μg of antibody in PBS (BSA-free) is mixed with 1 μl 0.33 mM CF750 Dye SE/TFP esters (Biotium 92142), Alexa Fluor 647 NHS Ester (Thermo Fisher Scientific A20006), Alexa Fluor 555 NHS Ester (Thermo Fisher Scientific A20009), or Alexa Fluor 488 NHS Ester (Thermo Fisher Scientific A20000) in DMSO, and incubated at room temperature for 16 hours. Fluorophore-conjugated antibodies were then purified with 30 kDa molecular weight cutoff Amicon Ultra-0.5 Centrifugal Filter Unit (Millipore Sigma UFC5030BK) by adding 500 μl PBS and centrifugation at 13,000 g for 5 minutes twice. Purified fluorophore-conjugated antibodies were collected by centrifugation at 2,000 g for 2 minutes and adjusted to 0.1 mg/ml in PBS for short-term storage at 4°C. Antibodies are listed in **Supplementary Table 5**.

### CRISPRmap optical pooled CRISPR knockout screen

HT1080/Cas9 AAVS1 cells were seeded into 6-well tissue culture-treated plates at a density of 50,000 cells/cm^2^. After 24 hours, cells were transduced with the GFP-pilot lentiviral supernatant supplemented with 8 μg/ml polybrene (Sigma-Aldrich TR-1003-G) at MOI ∼0.1. At 48 hours post-infection, viral supernatant was removed and cells were treated with media containing 2 μg/ml puromycin (Thermo Fisher Scientific A1113802) for 48 hours and seeded onto 96-well glass bottom plates (Cellvis P96-1.5H-N) at 10,000 cells per well. A total reaction volume of 50 μl was used in the following steps unless otherwise specified. After 24 hours, cells were fixed in 4% Paraformaldehyde (Electron Microscopy Sciences 15710-S) in PBS (Gibco 10010049) for 10 minutes at room temperature, followed by two rinses in PBS. Cells were then incubated in 0.1 mg/ml Wheat Germ Agglutinin (WGA) CF770 conjugate (Biotium 29059) and 0.5 μg/ml DAPI (Abcam ab285390) in PBS for 30 minutes at room temperature and imaged in PBS for membrane, GFP, and nuclei signal using the microscope configuration described above. After phenotype imaging, cells were permeabilized with 0.2% Triton X-100 (Sigma-Aldrich T8787) in PBS for 10 minutes at room temperature, followed by two rinses in PBS. For primer and padlock oligo hybridization, cells in each well were incubated in the Hybridization mix (GFP-pilot CRISPRmap Padlock and Primer mix (see **Supplementary Table 4** for sequences, each oligo in the mix has a final concentration of 10nM), 1 mg/ml yeast tRNA (Invitrogen 15401011), 2x SSC, 20% formamide (v/v) in ultrapure water) for 16 hours at 40°C in a HybEZ oven (ACD PN 321720). After hybridization, cells were first rinsed three times with the hybridization wash buffer (2x SSC, 20% formamide (v/v) in ultrapure water), then washed three times in the hybridization wash buffer for 5 minutes at 40°C. Cells were then incubated in splint mix (10nM CRISPRmap GFP-pilot splint mix (see **Supplementary Table 4** for sequences, each splint oligo in the mix has a final concentration of 10nM), 0.1% yeast tRNA, 2x SSC and 15% formamide in ultrapure water) for 30 minutes at 37°C in a HybEZ oven, rinsed twice with the formamide wash buffer (2x SSC, 15% (v/v) formamide in ultrapure water), and incubated in 2x SSC in ultrapure water for 15 minutes at room temperature. For T4 DNA ligation, cells were incubated in ligation mix (1x T4 ligase buffer, 1% (v/v) T4 DNA ligase (Enzymatics L6030-HC-L) in ultrapure water) for 2 hours at 16°C then 1 hour at 25°C in a HybEZ oven, followed by two rinses in PBS. For rolling circle amplification (RCA), cells were incubated in RCA mix (1x QualiPhi buffer, 2% (v/v) QualiPhi DNA Polymerase (4basebio 510100), dNTP mix, 0.25 mM each (Thermo R1122), 0.02 mM 5-(3-Aminoallyl)-dUTP (Thermo AM8439) in ultrapure water) for 6 hours at 30°C then remove the RCA mix and immediately fix with 4% PFA in PBS for 10 minutes at room temperature, followed by three PBS washes. For readout probe hybridization, cells in each well were incubated in readout probe mix (10nM of each readout probe (see readout probe sequences for each hybridization rounds in **Supplementary Table 4**), 2x SSC, 15% formamide in ultrapure water) for 30 minutes at 37°C in HybEZ oven. Cells were then imaged in the imaging buffer (0.5 μg/ml DAPI, 10 μg/ml Fungin (InvivoGen ant-fn-1) in PBS) using the microscope configuration described above. After imaging, the cells were incubated in the stripping buffer (2x SSC, 50% formamide (v/v) in ultrapure water) for 20 minutes at 40°C in the HybEZ oven and then rinsed once with formamide wash buffer. A total of 4 readout hybridization rounds were performed to decode all the CRISPRmap barcodes in the GFP-pilot library.

### CRISPRmap multi-omic optical pooled base-editing screen

MCF7-BE3 cells were transduced with the DDR364 library in the same manner as in the GFP-targeting knockout screen with several modifications to accommodate the multiplexed immunofluorescence and RNAmap. Specifically, after puromycin selection, cells were seeded onto 6-well glass bottom plates (Cellvis P6-1.5H-N) at a density of 50,000 cells/cm2. For the DDR364-irradiation screening, after 48 hours, cells were exposed to 10 Gy ionizing radiation using the Gammacell 40 cesium source irradiator and fixed at 6 hours after irradiation. For the DNA damaging agents (DDR364-chemo) screening, cells were treated with 100 nM Camptothecin (Sigma-Aldrich C9911), 1 μM Olaparib (Selleck Chemicals S1060), 1 μM Cisplatin (Sigma-Aldrich P4394), 1μM Etoposide (Sigma Aldrich E1383), or untreated, and fixed at 24 hours post-treatment. After fixation, cells were permeabilized with 0.1% Triton-X100 in PBS for 10 minutes on ice. Cells in each well were incubated in 1 ml reaction mix or buffers in all steps unless otherwise specified. After permeabilization, cells were incubated in the antibody mix (2 μg/ml rat anti-CD326 (BioLegend 312502), 1 μg/ml rabbit anti-RAD51-AF647 (Bioacademia 70-012), 2 μg/ml mouse anti-BRCA1-AF555 (Santa Cruz Biotechnology sc-6954), 0.5 μg/ml rabbit anti-RPA2-AF488 (Bethyl Laboratories A300-244A) in PBS, primary antibodies were fluorophore conjugated as described above) for 1 hour at room temperature, then rinsed twice with PBS. Cells were incubated in 10 μg/ml goat anti-Rat-IgG secondary antibody (Thermo Fisher Scientific SA5-10023) for 30 minutes at room temperature, and rinsed twice with PBS. Cells were fixed in 4% PFA in PBS for 10 minutes at room temperature to cross-link the antibodies to the cells, followed by two PBS rinses. Cells were then processed with padlock and primer probes hybridization, splint hybridization, ligation and RCA as described above, with the minor difference that 3nM of each CRISPRmap Padlock and Primer probes and 3nM of each RNAmap Padlock and Primer probes were used in the hybridization mix. Probe sequences are listed in **Supplementary Table 5.** After RCA and fixation, cells were first imaged in the imaging buffer for membrane, nuclei, and nuclear foci signal using the microscope configuration described in **Supplementary Table 5**. After imaging, the antibody signal was bleached with 1mg/ml lithium borohydride (Sigma-Aldrich 222356) and rinsed twice with PBS, prior to the incubation of the next round of antibodies. For both the DDR364-irradiation and the DDR364-chemo screening, a total of 4 antibody incubation-bleaching rounds were performed, the antibodies and conjugated fluorophores are listed in **Supplementary Table 5**. After the last round of bleaching, eight rounds of RNAmap readout probe hybridization-stripping rounds were performed, followed by eight rounds of CRISPRmap readout probe hybridization-stripping rounds. Each round was imaged using the microscope configuration described above. Readout probe sequences and conjugated fluorophores are listed in **Supplementary Table 5.** For the DDR364-irradiation screening, cells were incubated in Vector TrueVIEW Autofluorescence Quenching reagent (Vector Laboratories SP-8400-15) for 5 minutes at room temperature to reduce autofluorescence, followed by three rinses in PBS before imaging each CRISPRmap readout round in high DAPI imaging buffer (2.5 μg/ml DAPI, 10 μg/ml Fungin in PBS).

### In vivo CRISPRmap barcode readout and multi-omic phenotyping

Following lentiviral transduction with pLV-EF1a-TagBFP2 on OE19 cells, fluorescence-activated cell sorting (Sony MA900) was performed to obtain a BFP-expressing OE19 population. BFP-expressing OE19 (referred as OE19-BFP) cells were lentivirally transduced with the DDR364 library and puromycin selected as described above, then expanded for 4 days in puromycin-free media. We suspended 5×10^6^ cells in a 1:1 mixture of Matrigel and PBS and inoculated the mixture into the flanks of nude mice (JAX, strain no.002019). After 17 days, tumors were harvested and fresh frozen in OCT on dry ice and stored at −80°C. Frozen tumor samples were sectioned in Cryostat Microtome (Leica CM1510S) at −20°C into 10 μm thick sections, and deposited onto 12-well glass-bottom plates (Cellvis P12-1.5H-N) coated with 0.1 mg/ml poly-D-lysine (Sigma-Aldrich A-003-E). CRISPRmap barcode readout and antibody staining were performed as described above in the DDR364-irradiation base-editing screen on MCF7-BE3 cells with minor modifications. Specifically, 400 μl of reaction mix and buffers were added to each well to fully cover the tissue section. Tissue sections were fixed with 4% PFA in PBS for 15 minutes at room temperature and permeabilized with 0.5% Triton X-100 in PBS for 15 minutes at room temperature. 30nM of each CRISPRmap padlock and primer oligos were used in the hybridization mix. The same set of CRISPRmap padlock and primer probes, splints, and readout probes was used as in the DDR364-irradiation screening. Eight CRISPRmap readout cycles were performed prior to antibody staining and bleaching cycles. The same readout probes were used as in the base-editing screens. The antibodies and conjugated fluorophores are listed in **Supplementary Table 5**.

### Animal studies

Adult NU/J mice (>6 weeks of age) were obtained from The Jack Laboratory (Bar Harbor, ME). Animal care and experimental procedures were performed in accordance with the Guide for the Care and Use of Laboratory Animals of the National Institutes of Health and were approved by the Columbia University Institutional Animal Care and Use Committee (protocol AC-AABT8654). Mice were housed with a constant temperature of 21–24°C, 45–65% humidity and a 12-h light–dark cycle. All experiments were performed on female NU/J mice.

### Variant annotation

The sgRNA category of each guide was annotated as previously described^5^. We grouped the splice-donor and splice-acceptor categories into a ‘splice’ category. All AAVS1-targeting and non-targeting guides are annotated as a ‘control’ category. The ClinVar category was determined by querying each guide in the ClinVar database (v. 2023-12-15; https://www.ncbi.nlm.nih.gov/clinvar/; RRID:SCR_006169). Nonsense and missense variants were queried based on the specific amino acid change, whereas splice variants were queried based on the nucleotide change outcomes in the editing window (base C in the 4th to 8th bases in the sgRNA targeting sequences). Note that if multiple C bases exist in the editing window, a splice guide can render other mutational outcomes, such as missense or intron variants. These mutational outcomes were not counted in the annotation of splice variants but listed as “Less deleterious variants’’ in **Supplementary Table 5**. The determining criteria of the Clinvar category were established as previously described^5^. Briefly, three categories were assigned to non-control guides: i) benign/likely-benign (B/LB); ii) variants of uncertain significance (VUS); iii) pathogenic/likely-pathogenic (P/LP). The VUS category also includes variants with conflicting interpretations. If a variant was not documented in the ClinVar database, it was listed as “unknown”. All control guides were annotated as “control”.

### Library quality-check and quantification

For oligo pool quantification, the first-round amplification product in the library cloning step was collected and 0.5 μl of 50 μl PCR product was added to each 50 μl Q5 reaction mix for the second-round amplification of 10 cycles using the primer pairs CRISPRmap-F-ad and CRISPRmap-R-ad in **Supplementary Table 6**. We amplified 10 pg plasmid extraction product from the library cloning step with the same two-round strategy as the oligo pool quantification for plasmid-level quantification. Genomic DNA of the cells transduced with the sgRNA library were extracted with Genomic DNA Clean & Concentrator (Zymo Research D4010). We amplified 100 ng genomic DNA with the same two-round strategy. We had 5 ng of the final PCR product sequenced with next generation sequencing (Azenta Life Sciences Amplicon-EZ). Sequencing reads were analyzed to quantify the relative abundance of each guide in the library and the barcode-sgRNA recombination rate.

### Immunoblotting

We ordered individual sgRNAs as synthesized double-strand DNA fragments (integrated DNA Technologies) and cloned them individually into the CRISPRmap-CROPseq vector. Cells transduced with individual sgRNAs were selected for 2 days in puromycin and cultured for 2 days before collection as described in the base-editing screening. Cells treated with siRNAs were subjected to reverse siRNA transfection utilizing of firefly (FF) siRNA, BRCA1 siRNA, or BRCA2 siRNA at 20 nM and lipofectamine RNAiMAX (Thermo Fisher Scientific 13778075) as per manufacturer’s indications. Cells were trypsinized, washed and resuspended in sample buffer (0.1M Tris pH 6.8, 4% SDS, 12% β-mercaptoethanol) at a density of 20,000 cells/μl. Subsequently, samples were sonicated for 10 seconds twice and boiled at 95°C for 5 min prior to gel electrophoresis. After gel electrophoresis, proteins were transferred onto nitrocellulose membranes. Proteins were detected using the appropriate primary and HRP-conjugated secondary antibodies at a 1:10,000 dilution. Primary antibodies used in this study include mouse-anti-BRCA1 (Santa Cruz Biotechnology sc-6954, 1:100), rabbit anti-phospho-KAP1 (Bethyl Laboratories A700-013, 1:1,000), rat anti-Tubulin (Novus Biologicals NB 600-506, 1:50,000), and mouse anti-BRCA2 (Millipore OP95, 1:1,000).

### RNA sequencing

Gamma-irradiated and untreated MCF7-BE3 cells were prepared in parallel to the cells profiled in the DDR364-irradiation screen. Six hours after irradiation, total RNA was extracted with Quick-RNA Microprep Kit (Zymo Research R1051), and mRNA was isolated with NEBNext Poly(A) mRNA Magnetic Isolation Module (New England BioLabs E7490L). RNA integrity number (RIN) was quantified with RNA Pico 6000 assay (Aligent 5067-1513) on BioAnalyzer (Aligent 2100 G2939BA). DNA libraries for Next Generation Sequencing were prepared with NEBNext Ultra II RNA Library Prep Kit for Illumina (New England BioLabs E7775) and NEBNext Multiplex Oligos for Illumina (Unique Dual Index UMI Adaptors RNA Set 1) (New England BioLabs E7416). DNA libraries were quality checked with DNA 1000 assay (Aligent 5067-1504) on Bioanalyzer, then sequenced on a Miseq platform (Illumina) with a 5% spike-in of PhiX (Azenta Life Sciences Sequencing-Only). Four replicates were sequenced and the average Transcript per Million reads (TPM) was calculated for the transcripts we profiled optically with RNAmap.

### RNAmap primer and padlock probe design

The gene-specific target probes for RNAmap are designed for specificity, and minimized off-target binding, conforming to SeqFISH methodologies^19,31^. Utilizing the FISHprobe R package (v0.4.1; https://github.com/stevexniu/fishprobe), we established a systematic and reproducible protocol. For Gene Selection and Probe Extraction, we selected highly expressed gene isoforms from the Human GTEX V8^32^ and Mouse ENCODE^33^ tissue expression datasets for probe design. Probe sequences, 20-30 nucleotides in length, were derived from the coding sequence (CDS) and, where necessary, from the untranslated regions (UTRs). We targeted a GC content range of 45%-65% or 30%-70% for the targeting probes, excluding those with unsuitable GC content or sequences prone to forming homopolymeric runs (such as G-quadruplexes) to maintain optimal hybridization characteristics.

Specificity and Off-Target Minimization: Local BLAST^34^ searches against human and mouse mRNA sequence databases identified probes with off-targets, particularly those with alignments exceeding 10-15 nucleotides with unrelated genes in the transcriptomes and the repetitive DNA using repetitive masks. Tissue-specific expression data from human^32^ or mouse^33^ were pivotal in developing a gene copy-number table for each tissue type, which informed the exclusion of probes with off-target copy numbers exceeding 15-20 logTPM. For thermal stability and structural integrity, to refine the probe pool by optimizing GC content for enhanced binding affinity, an iterative selection process was employed. Probes were initially ranked in ascending order of their deviation from the target GC content of 55%, starting with the probe exhibiting the greatest deviation. This arrangement continued until no overlapping probes remained. Subsequently, the selection process took into account the calculated melting temperatures (Tm)^35^. For secondary structure predictions, including pseudoknots, the analysis was conducted under specific conditions: a sodium ion concentration of 0.33 M (equivalent to 2X SSC), and 50% formamide at 37°C^35^. Probes with an equilibrium stability below 20% were excluded to ensure the formation of stable and specific duplexes. For final probe set selection, the finalized probe set, consisting of 28-32 probes per gene, was optimized to minimize spatial overlap, allowing a maximum of 5 nucleotides of overlap between adjacent probes. Probes were subjected to stringent filters for equilibrium, and free energy, to refine the probe library. Local BLAST searches within the probe pool identified and mitigated potential cross-hybridizations between the selected probes. Genes with insufficient probe numbers were curated manually using a genome browser to guarantee thorough coverage. For isoform-specific probes, in parallel to the above strategy, isoform-specific probes were selected based on the unique exon-intron structures of the isoforms. These probes were designed to span exclusive exon-junction regions to ensure specificity to distinct transcript variants. A refined BLAST strategy, specialized for isoform differentiation, focused on these unique splice junctions, allowing us to curate probe sets with sequences that exclusively mapped to the target regions.

### CRISPRmap and RNAmap readout probe, CRISPRmap padlock and primer oligo design

For probe generation and off-target screening, starting with a base of 240,000 25-mer probes 8, we generated all possible 20-mer sequences sequentially. Each of these subsequences was subjected to BLAST screening against human and mouse transcriptomes to exclude any probes with off-target complementarity, and the resulting pool was thus reduced to only those probes with zero off-target hits. For optimizing Probe Performance: To optimize the readout probes’ performance, we calculated their melting temperatures (Tm) and secondary structure predictions^35^ to refine our selection further similar to the aforementioned target probes. This ensures that each probe binds to its intended target with high affinity and that the thermal profiles are suitable for our experimental conditions. In scenarios with high mRNA expression, it is vital to prevent overcrowding within any single fluorescence channel. Additionally, based on expression levels in the targeting tissue (GTEX^32^ and ENCODE^33^) we curated the probe sets and their corresponding fluorophores, and distributed the signal across multiple channels, promoting distinct visualization of each mRNA molecule. To further minimize the risk of cross-hybridization, we conducted an analysis of readout probe sequences for potential overlaps by performing a local BLAST search against the readout probe pool. This effort led to the identification of 226 20-mer DNA sequences, as outlined in **Supplementary Table 6**, which provides details for each probe, including the 20-nt probe sequence, unique identifier, off-target information, melting temperatures (Tm), and secondary structures. For codebook construction: By employing a Hamming distance approach, similar to the HDM4 code used in MERFISH^14^, a codebook was constructed with 36 of the aforementioned 226 20-mer readout probes. This codebook consists of 319 36-bit codes allocated over 12 hybridization rounds across three channels (488nm, 561nm, 640nm), ensuring that each readout probe would have a unique signature, reducing the possibility of channel crosstalk and fluorescence overlap. This approach aims to enable differentiation of probes even in densely labeled samples, where multiple mRNA molecules are in close proximity. The detailed codebook design is provided in **Supplementary Table 6,** which includes details such as binary code assignments for each hybridization round and optical channel, indices, a conversion table that relates binary codes to specific probes, and sequences linked to each code across channels. For CRISPRmap readout probes, we selected 24 20-mers from the aforementioned 226 20-mer list. This selected set of 24 probes was split into 2 sets of 12 probes for the detection of Padlocks and Primers, respectively. Splints sequences consist of the reverse complement of the primer readout sequences, with an additional universal 2 bases added at the 5-prime end (“GT”), and the 3-prime end (“AC”) in an attempt to avoid ligation efficiency biases between different splint oligos. Two sets of 54 30-mer encoding sequences were generated with similar criteria as the 20mer list, and used as Padlock or Primer encoding sequences. The sequences of these oligos are listed in **Supplementary Table 6**.

### Image processing and analysis

#### Image storage and Stitching

All microscopy images were acquired using Fusion software and saved as ims files. Each ims file stores the image as a 5-D object in the order: Resolution, Channel, Z, Y, X. All image montages were stitched using Fusion’s stitching software. For 60x images, high-speed setting was used to stitch the image montage and saved as ims files. For the 20x images the high-quality setting using default parameters was used to stitch the image montage and saved as ims files.

### Image rescaling

All images were uploaded to a google cloud virtual machine for further image processing and analysis. Images were processed using a jupyter notebook (python). To read the ims file and the corresponding metadata we used “imaris_ims_file_reader” package. All 60x montages were analyzed at resolution 3 (1/8 scale of original image), and for all 20x images, we used resolution 1 images (1/2 scale of original image). All images were max projected along the Z-axis. All max projected images are 3 dimensional with the dimensions being Channel, Y, X. The images are in numpy array format and of uint16 data type. Our imaging protocol involves imaging cells at different magnifications based on the resolution of images required. If a particular imaging round was imaged at a different magnification, the images of this round will be of a different size and have a different pixel pitch (pixel to micron ratio). To accommodate for this, images were scaled to achieve a consistent pixel pitch using cv2 resize with a bicubic interpolation function. This also ensured that images from all imaging rounds were the same dimensions across X and Y.

### Image registration

After rescaling images across the multiplexed imaging rounds, we register images to a reference image. Across the multiplexed imaging rounds there are global translational shifts, (i.e. misalignment of the glass bottom well plate), as well as local translational shifts (i.e. cells slightly shifting between imaging rounds). To finely align the images across all imaging rounds we calculated the transformation matrices for each round using the TV-L1^16^ implementation of optical flow on binary nuclei masks derived from DAPI stains. Optical flow calculates Y, X vector shifts across the images for every pixel and performs pixel level registration. The transformation matrix was then applied to all image channels of that imaging cycle. During registration all images are converted from uint16 to float64. The images are then converted back to uint16 to reduce the memory usage and speed up image processing. All registered images are 3 dimensional with the dimensions being Channel, Y, X. Registration quality was estimated using cross-correlation. It is expected that the cross-correlation would decrease with increasing montage size. For our 30×30 60x montages and 10×10 20x montages a cross-correlation of > 0.75 was considered good.

### Segmentation of cell and nuclear boundaries

To assign detected guides to each cell and quantify nuclear antibody stains, we segmented both the cytoplasmic area and nuclear area of each cell using Cellpose^32^. This process was broken into three steps, pre-processing, segmentation, and filtering. The EPCAM and DAPI stains were preprocessed by thresholding to maximize the dynamic range of the plasma membrane and nuclear stains. Typically, this involved setting pixels below the 2nd percentile to 0 and pixels above the 98th percentile to 255. Following pre-processing, images were segmented twice with Cellpose, first to identify cytoplasmic areas and second to identify nuclear areas. The cytoplasmic segmentation run was performed on an image stack containing the EPCAM and DAPI stain and excluded any cytoplasm mask smaller than 5 pixels, while the nuclear segmentation was only performed on the DAPI stain with no minimum size requirement. The cell diameter parameter for Cellpose was determined by hand counting and averaging the width and height of 10 randomly sampled cells in pixels. This value was multiplied 1.5x for the cytoplasmic segmentation, and 0.75x for nuclear segmentation.

Once the nuclear and cytoplasmic masks were generated, we filtered out nuclear masks which did not overlap with a cytoplasmic mask, and cytoplasmic masks which did not contain a nuclear mask. This ensured that each segmented cytoplasm had one associated segmented nucleus and vice versa. The coordinates for each nuclear and cytoplasmic pair were relabeled with the nuclear ID, which was used as the cell ID from this point onwards. Segmentation quality was validated by both quantifying the percent of proposed cytoplasmic masks retained after filtering for segmented nuclei overlap and visual inspection of the images.

### CRISPRmap and RNAmap amplicon detection

To detect amplicons corresponding to CRISPRmap, all registered images corresponding to CRISPRmap readout rounds were processed as follows. Each 2D image (Y and X) underwent contrast stretching to improve the signal to noise ratio using skimage rescale_intensity function. Images were now stored in a list in the order of the readout round and channel (R1-ch1, R1-ch2, R1-ch3, R2-ch1…) with R1 being the first readout round and ch1 being the longest wavelength channel. For each image (readout round and channel combo) spots were identified by using the skimage implementation of the difference of gaussians method using parameters that maximized the barcode recovery. This implementation outputs an array of coordinates of all spots identified. All the coordinates for spots identified were searched against the cell masks (from cell segmentation) and any spot outside cell masks were discarded from further analysis. Furthermore, if the number of spots within a cell mask was < 3 then the spots within the cell mask were also discarded from further analysis. This was done to reduce the noise/error in spot detection. All the spots retained for a given round-channel image were stored in an array. This was repeated for each cycle-channel image and the array of spots retained was stored in a list (with the order being spots for R1-ch1, R1-ch2, R1-ch3, R2-ch1…). Another array was created combining all the retained spots across all imaging round-channel combos. To eliminate duplicates in the merged array we used the np.unique function and discarded spots within a radius of 2 pixels. Then for each spot coordinate in the merged array, we compare the distance of the spot with all the spots detected in a single round-channel combo. If the spot is within a 2-pixel radius we mark the given round-channel combo as positive. This was done for all rounds and channels and by doing so for each spot a “spot code” was generated. A spot code essentially maps for a given spot, which round and channel combinations also contain that spot. Once the spot code is generated for all the spots, the spot code for every spot was compared to the predefined barcode designed for every guide. If a spot code matched a barcode, the spot was assigned to the barcode. If a spot did not map to any barcode corresponding to a guide, then the spot was discarded from further analysis. Spot calling was optimized to maximize spots that are assigned to barcodes of the guide library.

### Guide assignment to cell masks

Finally, each cell was assigned a barcode based on the spot identity underneath the cell masks. For each cell mask the maximum number of spots within the cell mask corresponding to a single guide was identified (g1). If the number was greater than 2 then, the second highest number of spots within the cell mask corresponding to a single guide was calculated (g2). The ratio of g1/g1+g2 was calculated and if the ratio was greater than 0.67 then the cell was assigned to the guide corresponding to the largest number of spots within the cell mask (g1). Cells with < 3 spots for any guide and purity < 0.67 were discarded from further analysis. The barcode identity of the cell was stored in a dictionary as well as in the format of an image mask.

### Foci and micronuclei detection and quantification of antibody stains

All the registered images for the multiplexed antibody stains were obtained. For each cell, the sum intensity underneath a given cell mask was calculated for a given antibody stain and stored in a dictionary. The sum intensity underneath the nuclear mask was also calculated and stored in the dictionary. Also, the average intensity of each antibody stain was also calculated by dividing the sum intensity by the total number of pixels underneath the cell/nuclear mask. The raw images then underwent contrast stretching using the rescale_intensity function of skimage. After rescaling the images, foci detection for RAD51, BRCA1, RPA2, γH2AX, 53BP1 and RAD18 was done using skimage difference of gaussians method. The total number of foci within the cell mask/ nuclear mask was also stored in the dictionary.

To detect and quantify the presence of micronuclei, for every cell mask in the image, the nuclear mask was retrieved. If there was a nuclear mask of < 100 pixels in area, then the nuclear mask was separately annotated as a small nucleus.

For micronuclei detection, we first perform nuclei segmentation on the DAPI stain using Cellpose. Nuclei masks were generated to define the outline of each nucleus in the image. All the nuclei with a size less than 100 pixels were marked as “micronuclei”. This threshold was determined by manually inspecting the micronuclei captured by the nuclei segmentation. We then subtract the DAPI signal underneath all nuclei masks by changing the intensity value to 0 for all pixels outlined by the nuclei masks. We remove the background DAPI signal by changing the intensity value to 0 for any pixel with an intensity value below 110. To completely remove the residual DAPI signal coming from the DAPI staining of the cell nuclei, we dilate each nuclei mask by 2 pixels using cv2 then changing the intensity value to 0 for all pixels outlined by the dilated nuclei masks. We finally perform spot calling to identify micronuclei. Based on the coordinates of the spots, the number of spots within a cell mask were identified and were included in the dictionary as the number of micronuclei within a cell mask.

For further analysis, the dictionary was converted into a CSV file. Each CSV file contained the following information: cell-id, nuc-id, Avg intensity of EPCAM under membrane, Avg intensity of γH2AX under nuclei, Avg Ki67 under Nuclei, Avg cycB1 under Membrane, Avg cycB1 under Nuclei, Avg phoH3 under Nuclei, Avg cycA under Nuclei, Avg cPARP under Nuclei, Avg p21 under Nuclei, Avg p53 under Membrane, Avg p53 under Nuclei, RAD51 spots, BRCA1 spots, RPA2 spots, γH2AX spots, 53BP1 spots, RAD18 spots, small nuclei, micronuclei, Nuclei size in pixels, Membrane size in pixels. The number of spots for the foci features, the average intensity of protein stains, and the nuclear and membrane pixel sizes for each cell that passed barcode QC in the DDR364-irradaition screening are listed in **Supplementary Table 7**.

### Bootstrapped Wasserstein Distance

We computed a bootstrapped Wasserstein distance to measure the foci formation deviation from perturbation to control guides. We denote X_{g},j_ ∈ Z^|g|^ and X_{ctrl},j_ ∈ Z^|ctrl|^ as cells undergoing a specific perturbation guide or control respectively, where |g|, |ctrl| refer to the corresponding number of cells in each condition. For each perturbation g_i_ ∈ [[G]] and feature j ∈ (RAD51, BRCA1, RPA2, γH2AX, X53BP1 and RAD18), we 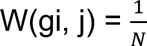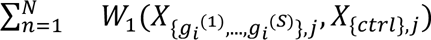 as the average 1-Wasserstein distance between guide i and control across S=50 samples with N=200 iterations, where in each iteration S cells {g_i_^(1)^,…, g_i_^(S)^}⊆ {g_i_} under guide i are randomly sampled without replacement. As a baseline control, we also computed 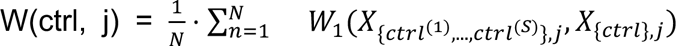 as the average 1-Wasserstein distance between randomly subsampled control cells and the full control cell set. The choice of bootstrapping cells is to mitigate the bias introduced by noticeable sample size differences across guides. We report the bootstrapped distances across perturbation guides and the control baseline with violin plots (**Supplementary Figure 4j-k**). Highlighted guides are chosen from the aforementioned significant hits with absolute log2-fold change (L2FC) > 0.5 and KS-test p-value (p.adj) < 0.05 under Benjamini-Hochberg correction.

### Beta-Binomial Test

We define the data as X ∈ Z^N^ ^×2^, where X_i_:= (n_i_, k_i_), where n_i_ denotes the number of cells affected by a given guide, g_i_, and k_i_ denotes the number of cells exceeding a fixed threshold for a cell-specific continuous or discrete feature. We refer to these cells as “positive” cells. Each guide, g_i_ ∈ [[G]], corresponds to either a control guide or a test guide defined by a mapping φ(g_i_) : [[G]] → {0, 1}, where φ(g_i_) = 1 corresponds to a test guide. To construct a plausible null hypothesis, we fit a Beta-Binomial distribution to the control data, X_ctrl_:= {X_i_ : φ(g_i_) = 0}. We use a Beta-Binomial distribution since we assume the number of positive cells is independently and identically distributed according to a Binomial distribution given the number of total cells for a guide. To account for overdispersion attributed to variability between control guides, we place a Beta distribution on the success probability of the Binomial distribution. For each condition the cells are placed in, we run a separate statistical test since we assume the rate of positive cells is significantly impacted by the environment, and we would like to test for the significance of specific guides conditioned on the environment.

The model is as follows:

p_i_ ∼ Beta(α, β)
k_i_ ∼ Binomial(n_i_, p_i_),

where the parameters α, β are inferred via a maximum likelihood estimation. Then, over the test data, X_test_:= {X_i_ : φ(g_i_) = 1}, we compute p-values according to a two-tailed test. These p-values are then adjusted using the Benjamini-Hochberg correction procedure to control for the false discovery rate (FDR). Finally, we reject the null for any corrected p-values falling below the significance level of 0.05. To ensure the null hypothesis is plausible with respect to the control data, we check to see if the computed p-values are uniformly distributed over the [0, 1] interval. The proportions, total number of cells, and adjusted p-values for each test guide are reported in **Supplementary Table 8**.

### Statistical analysis and graphic representation

Histograms were plotted with the histplot function in the seaborn package in python. Boxplots were plotted with the boxplot function, and violin plots with the violinplot function in the seaborn package in python. On the boxplots, the box shows the median, lower and upper quartiles while the whiskers extend to show the rest of the distribution, except for points that are determined to be “outliers” using a method that is a function of the inter-quartile range. Two-sided Mann Whitney U tests were performed to test the difference between the distributions using the statannotations package in python. *p < 0.05, **p < 0.01, ***p < 0.001, ***p < 0.0001. For hypothesis testing, the U test computes the exact p value by comparing the observed U statistic against the exact distribution of the U statistic under the null hypothesis. Scatter plots and volcano plots were plotted with the scatterplot function in the seaborn package in python. Foci number CDFs were plotted with the ecdfplot function in the seaborn package in python. The difference in foci number distribution was tested by a two-sided Kolmogorov–Smirnov test using the ks.test function in R and the p values were adjusted by the Benjamini-Hochberg method. *p.adj < 0.05, **p.adj < 0.01, ***p.adj < 0.001, ****p.adj < 0.0001. Guides with p.adj<0.05 and an absolute L2FC > 0.5 were regarded as statistically significant. One-sided Fisher exact test for gene enrichment was performed with the fisher.test function in R with the alternative hypothesis being “greater” and the p values were adjusted by the Benjamini-Hochberg method. Genes with an adjusted p-value < 0.05 were regarded as statistically significant. Heatmap for optical feature correlations and hierarchical clustering of guides were generated with the clustermap function in the seaborn package in python, method “complete” was used for clustering. Schematics were generated in BioRender.

## Data availability

All raw imaging data are being deposited to the Bioimage Archive (https://www.ebi.ac.uk/bioimage-archive/). Please reach out to the corresponding author for access.

## Code availability

All code is deposited on GitHub at https://github.com/GaublommeLab/CRISPRmap-Pipeline, and https://github.com/stevexniu/fishprobe

---

**Supplementary Figure 1.**
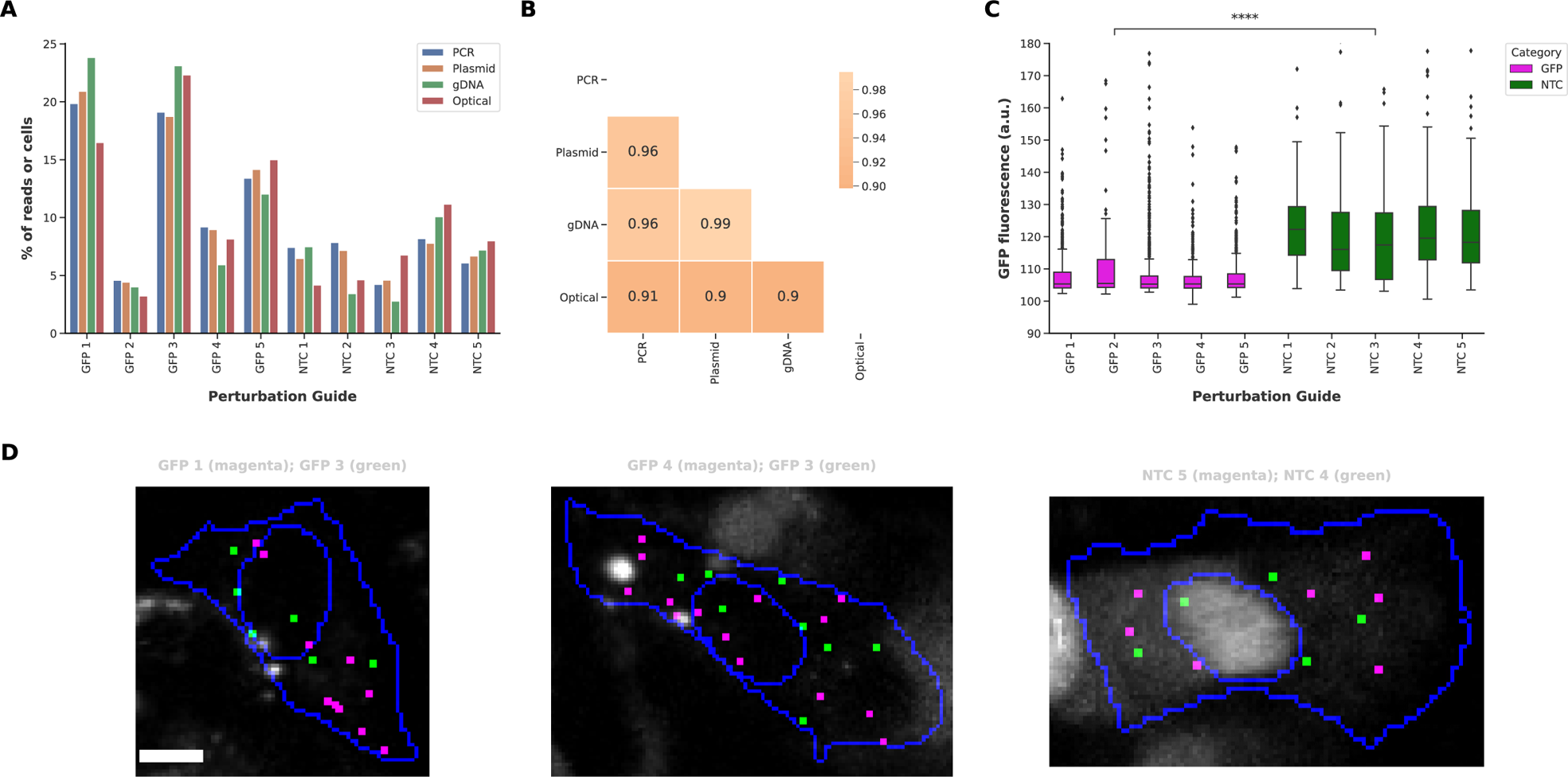
Performance of the GFP-targeting pooled optical CRISPR knockout screen. **(A)** Quantification of the relative abundance of guides in the pooled sgRNA library, showing good maintenance of guide representation in the pooled library based on the percentage of sequencing reads in PCR-amplified oligo pool (PCR), plasmid pool (Plasmid), and genomic DNA (gDNA) of cells transduced with the library, as well as the percentage of barcode-assigned cells in the optical screen (Optical). **(B)** Correlation of relative abundance of guides in the pooled sgRNA library at PCR, Plasmid, gDNA, and Optical level, showing high Pearson correlation above 0.9 between the PCR and Optical abundance. **(C)** Quantification of genotype-phenotype mapping, showing the cells with each GFP-targeting guide have significantly reduced GFP protein level compared to the cells with each non-targeting control guide. Pairwise test was performed on each GFP-NTC guide pair, and the GFP2-NTC3 pair with the least significant p-value is shown. Two-sided Mann-Whitney test, *p < 0.05, **p < 0.01, ***p < 0.001, *****p < 0.0001. **(D)** Visualization of double-transduced cells. Decoded barcodes reporting on the most (magenta) and second most (green) representing guides in each cell are shown as spots.Raw GFP fluorescence is displayed in greyscale. Cell and nuclear boundaries are outlined in blue. Scale bar, 20 µm.

**Supplementary Figure 2.**
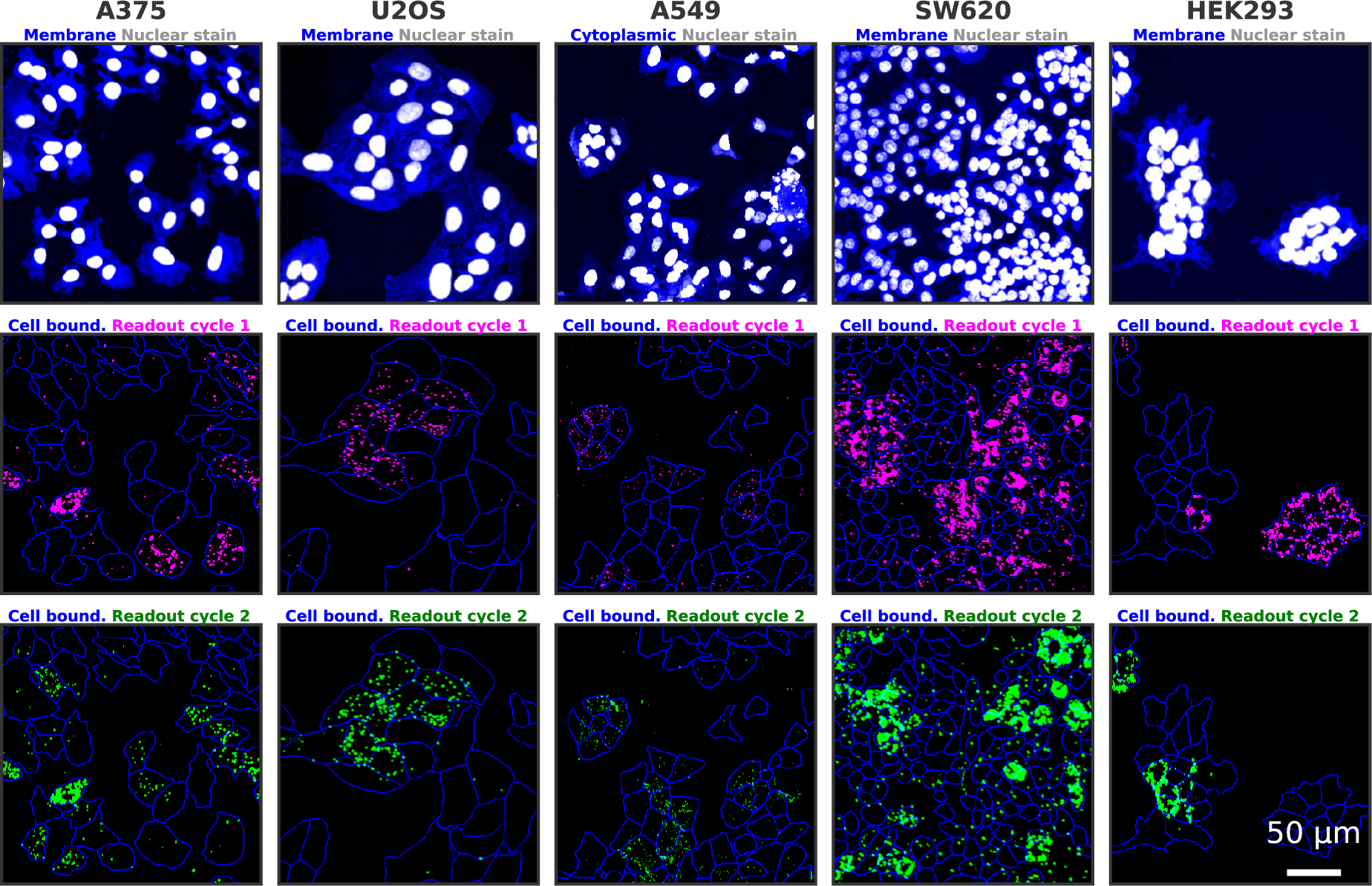
CRISPRmap application on multiple cell lines. Visualization of barcode detection on five cell lines: A735, U2OS, A549, SW620, and HEK293 (left to right). The morphology of each cell line is shown (top row) with membrane or cytoplasmic stains (blue) and nuclear stain (white). The fluorescence signal from two CRISPRmap imaging cycles are shown in magenta (middle row) and green (bottom row). Cell boundaries are outlined in blue. Scale bar, 50 µm.

**Supplementary Figure 3.**
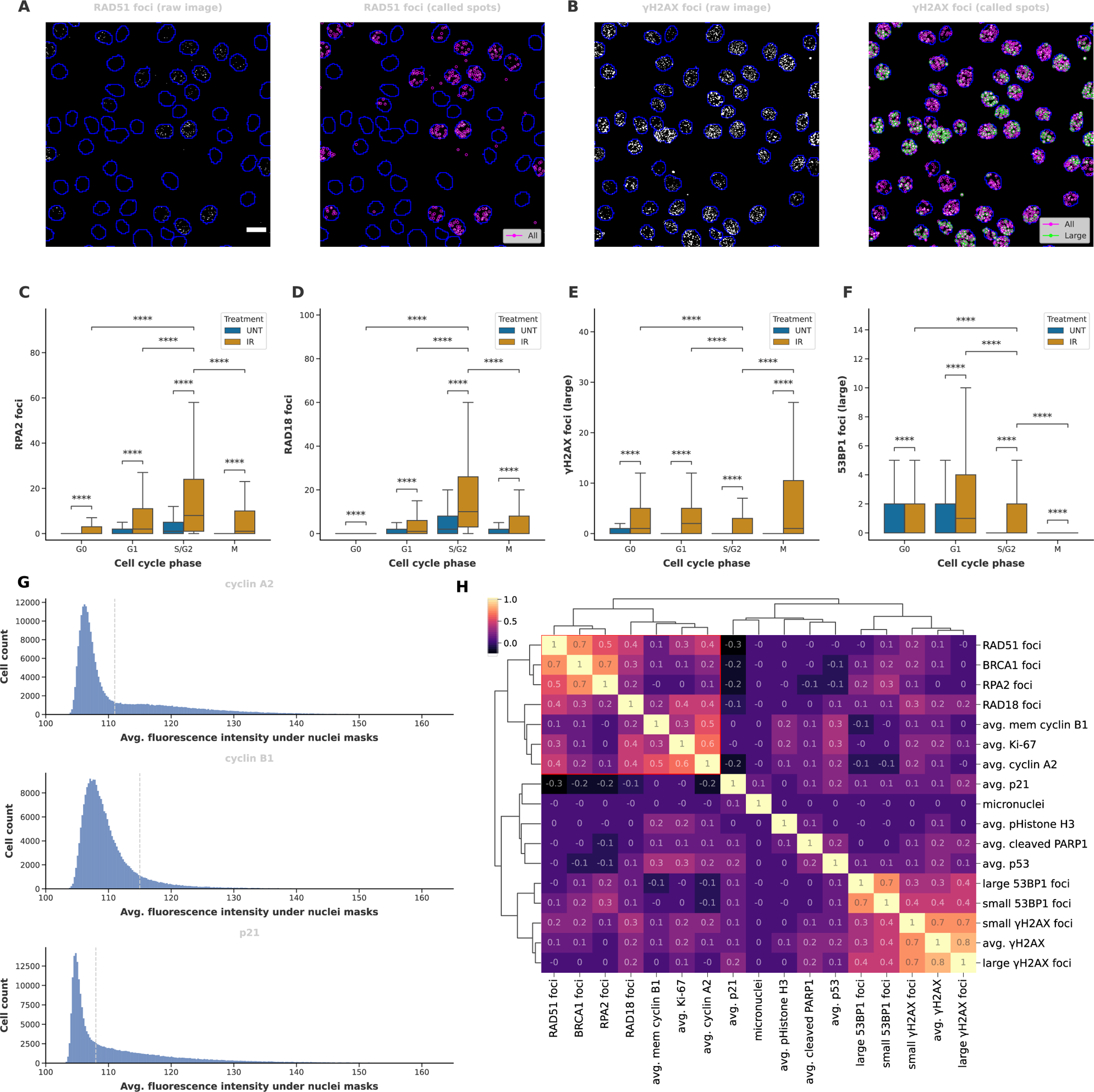
Multiplexed immunofluorescence enables cell cycle phase separation and nuclear foci detection. **(A)** Visualization of nuclear foci detection for RAD51. Raw antibody stain for RAD51 is shown in grayscale in both left and right panels. Nuclear boundaries outlined in blue. All computationally detected RAD51 foci are displayed as circles in false color (magenta) on the right panel. Scale bar, 20 µm. **(B)** Same as A) for γH2AX. All computationally detected γH2AX foci are displayed as circles in false color (magenta) on the right panel, whereas computationally detected spots categorized as large γH2AX foci are displayed as circles of false color green. The same region in A) is shown. **(C)** Quantification of the number of RPA2 foci per cell across cell cycle phases in untreated (UNT) and irradiated (IR) cells. Two-sided Mann-Whitney test, *p < 0.05, **p < 0.01, ***p < 0.001, ****p < 0.0001. (Outliers are omitted in the plot.) **(D)** Same as C) for RAD18 foci. **(E)** Same as C) for large γH2AX foci. **(F)** Same as C) for large 53BP1 foci. **(G)** Quantification of protein expression level of cyclin A2, cyclinB1, and p21. A threshold on the average fluorescence intensity under nuclear masks is set to categorize the expression level of the corresponding protein into low (smaller than the threshold) or high (greater than or equal to the threshold) categories. The threshold for each protein is displayed as a dashed line on the corresponding histogram. **(H)** Correlation of optical features measured in this experiment, showing positive correlation among nuclear foci involved in homologous recombination (RAD51, BRCA1, RPA2, RAD18), S/G2 phase cell cycle markers (cyclin A2, cyclin B1) and proliferation marker (Ki-67).

**Supplementary Figure 4.**
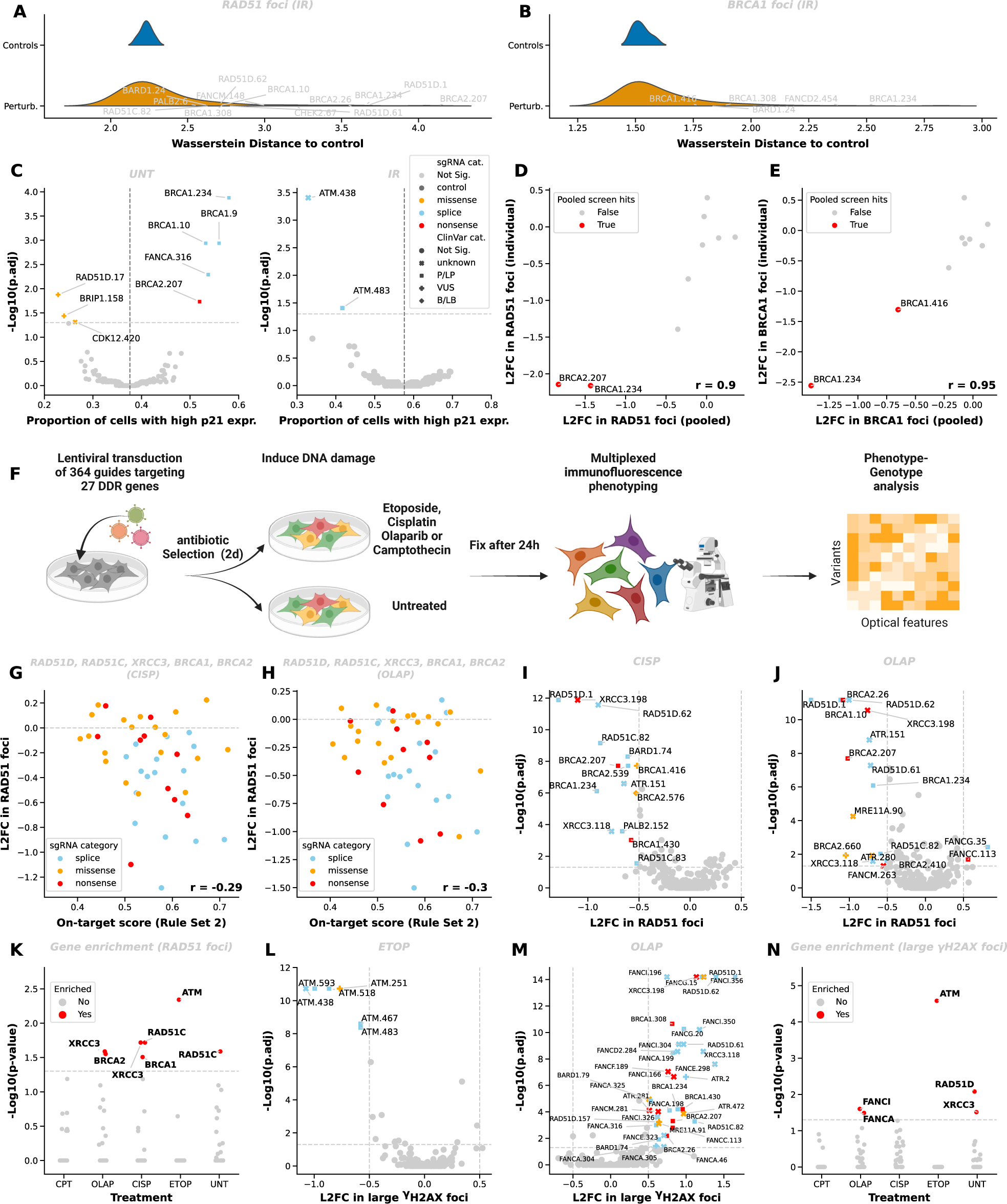
Performance of CRISPRmap base-editing screening on MCF7 cells treated with DNA damaging agents. **(A)** Wasserstein distance of cells with DDR gene-targeting guides (Perturb) or control guides (Controls) to control cells for RAD51 foci in irradiated cells. Hits of RAD51 foci identified in the pooled screening are marked. **(B)** same as A) for BRCA1 foci. **(C)** Volcano plots showing variants yielding significant changes in the proportion of cells with high p21 expression in untreated (left) or irradiated (right) conditions. Significance is defined by p.adj < 0.05, Beta-Binomial test. The proportion of control cells with high p21 is marked with the vertical line (UNT: 0.376; IR: 0.576). sgRNA category and ClinVar category for significant guides are displayed by colors and shapes, respectively. **(D)** Correlation of RAD51 foci log2-fold change (L2FC) in guides delivered in the pooled library (pooled) and transduced individually (individual). The Pearson correlation (r) is 0.90. **(E)** same as D) for BRCA1 foci. The Pearson correlation (r) is 0.95. **(F)** MCF7-BE3 cells were lentivirally transduced with the DDR364 library, selected in puromycin-containing medium for 2 days, and cultured 2 days without puromycin. Cells were either untreated (UNT) or treated with DNA damaging agents Camptothecin (CPT), Olaparib (OLAP), Cisplatin (CISP) or Etoposide (ETOP) for 24 hours, then fixed before optical phenotyping and barcode detection. **(G)** Correlation between L2FC in RAD51 foci in CISP-treated cells and the Rule Set 2 on-target score, showing a negative correlation. All guides targeting RAD51 regulators including RAD51D, RAD51C, XRCC3, BRCA1, and BRCA2 are shown. The Pearson correlation (r) equals −0.29. **(H)** same as G) for OLAP-treated cells. The Pearson correlation (r) equals −0.30. **(I)** Volcano plot showing variants yielding significant changes in RAD51 foci in CISP-treated cells. Statistical significance is defined by p.adj < 0.05 and absolute L2FC > 0.5. All guides targeting DDR genes with on-target score >= 0.5, all AAVS1-targeting and non-targeting control (NTC) guides are shown. Two-sided KS test, *p.adj < 0.05, **p.adj < 0.01, ***p.adj < 0.001, ****p.adj < 0.0001. Same figure legend as in C). **(J)** same as I) for OLAP-treated cells. **(K)** Gene enrichment analysis in variants that result in significant changes in RAD51 foci. Fisher exact test. Enrichment is defined as p.adj <0.05. **(L)** Same as I) for variants yielding significant changes in large γH2AX foci in CISP-treated cells. **(M)** same as L) for OLAP-treated cells. **(N)** same as K) for large γH2AX foci.

**Supplementary Figure 5.**
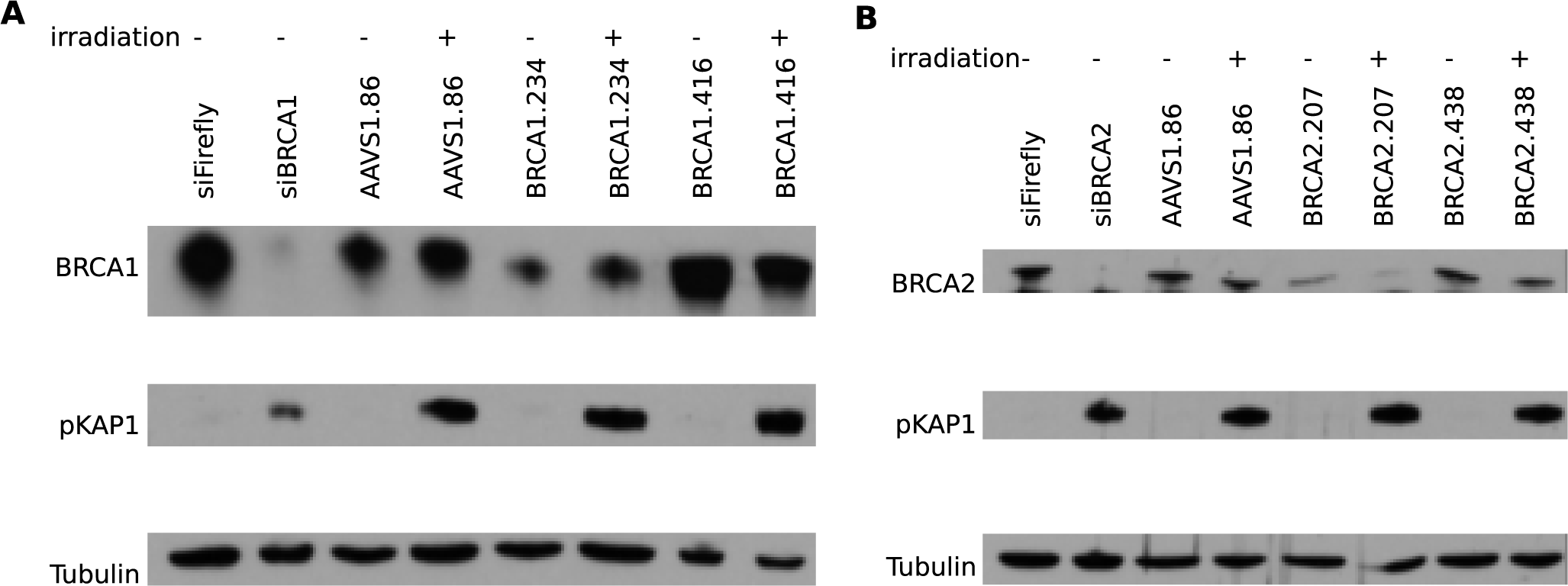
Immunoblots on cells with individually transduced guides. **(A)** Immunoblots on guides targeting BRCA1 showing the reduction of full-length BRCA1 protein in the splice variant BRCA1.234, compared to the missense variants BRCA1.416 and.BRCA1.476, and the AAVS1 variant AAVS1.86. Cells transduced with Firefly siRNA (siFirefly) and BRCA1 siRNA (siBRCA1) were included to show the specificity of BRCA1 detection. Cells were treated with or without irradiation and the induction of DNA damage is shown with the phospho-KAP1 (pKAP1) staining. Tubulin is used as the loading control. **(B)** As in A) for BRCA2 variants showing the reduction of full-length BRCA2 protein in the nonsense variant BRCA2.207, compared to the missense variant BRCA2.438 and the AAVS1 variant AAVS1.86.

**Supplementary Figure 6.**
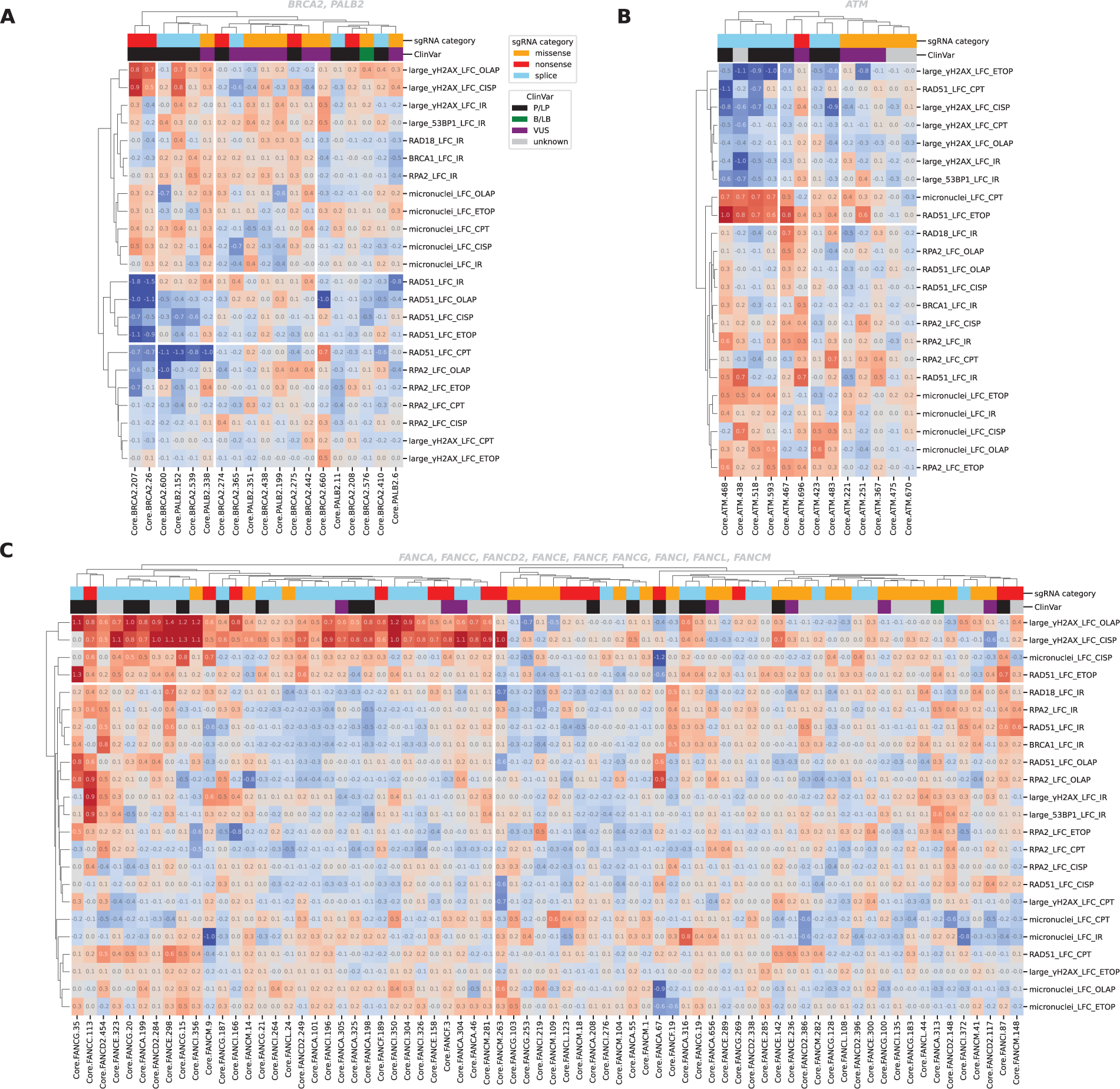
Variant analysis of functionally relevant genes and variant clusters with treatment-specific optical signatures. **(A)** Clustering of guides targeting BRCA2 and PALB2, showing a cluster of pathogenic nonsense variants showing a string reduction of RAD51 across all treatments and another cluster showing reduction of RAD51 in CPT-, OLAP-, and CISP-treated cells mainly composed of pathogenic splice variants. Other clusters show mild phenotypes. Log2-fold change (LFC) in each optical phenotype in corresponding treatment conditions are shown as rows in the heatmap. Cells in all cell cycle phases are included, untreated cells are not included. All guides with on-target score >= 0.5 were included in the clustering and shown in the heatmap. Columns were cut at a depth of 2 and rows were cut a depth of 3 based on the dendrogram. Values of L2FC are displayed on the heatmap. Color scale is −1 to 1. **(B)** same as A) for guides targeting ATM, showing a cluster with reduced large γH2AX foci and micronuclei in all treatments, increased micronuclei in CPT-treated cells and increased RAD51 foci in ETOP-treated cells composed of splice variants, and other clusters with milder phenotypes. **(C)** same as A) for guides targeting all genes in Fanconi anemia (FA) pathway, showing a cluster with increased large γH2AX foci in OLAP- and CISP-treated cells, increased micronuclei in ETOP-treated cells and increased RAD51 foci in ETOP-treated cells, composed of most splice and nonsense variants targeting these genes. The other cluster composed of mainly missense variants shows mild phenotypes.

## Supplementary Table information

**Supplementary Table 1. Cost estimation of the CRISPRmap assay.** The cost of probes and fluorophores, key chemicals and reagents, and enzymatic reactions are listed. The calculation of cost per million cells is based on the base-editing screening on MCF7 cells using the DDR364 library. The actual cost may differ according to price changes on the vendors’ side. The total setup cost mainly results from the purchase of Padlock, primer, splint and readout probes, which require a minimum amount to be ordered.

**Supplementary Table 2. GFP-pilot data analysis.** This table shows the cells analyzed in the CRISPRmap knockout screening targeting GFP, related to Figure 2 and Supplementary Figure 1. The number of amplicons detected as well as the average nuclear GFP intensity in each cell are listed. Cells that passed QC are listed with their unique guide identities. The pairwise Mann-Whitney U test p values of each GFP-NTC guide pair are listed, related to Supplementary Figure 1c.

**Supplementary Table 3. OE19 tissue barcode readout data analysis.** This table lists the cells analyzed in the OE19 tissue profiling, related to Figure 6. The number of decoded barcodes for the top two most abundant guides, the guide purity, and the QC status are listed for each cell.

**Supplementary Table 4. GFP-pilot library design and readout scheme.** This table lists the sgRNA_ID, sgRNA sequences and barcode sequences in the GFP-pilot library, as well as the sequences of padlocks, primers, splints and readout probes that were used to readout the barcodes, related to Figure 2 and Supplementary Figure 1. A detailed readout scheme including the order, conjugated fluorophores and imaging setting of the 8 readout probes are listed, as well as the 8-bit codebook encoding the 10 guides in the library.

**Supplementary Table 5. DDR364 library design, readout scheme, and antibody staining.** This table lists the sgRNA_ID, sgRNA sequences and barcode sequences in the DDR364 library, related to Figure 3, 4, 5 and Supplementary Figure 3, 4, 5. The sequences of padlocks, primers, splints and readout probes that were used to readout the barcodes and a detailed readout scheme including the order, conjugated fluorophores and imaging setting of the 24 readout probes are shown, together with the 24-bit codebook encoding the 364 guides in the library. The mutational outcomes and ClinVar annotations of the guides are listed. Antibodies used in the irradiation screen, DNA damaging agents screen (chemo) are listed in detail. The antibodies used in OE19 tissue profiling are also listed in detail, related to Figure 6.

**Supplementary Table 6. Oligonucleotide sequences of CRISPRmap probes.** This table lists the sequence of all the 54 padlocks and 54 primers used in the DDR364 base-editing screening. All 2,916 possible padlock-primer combinations are listed with their corresponding readout sequences specified. This table also lists the sequences of 12 splints and 24 readout probes that are required for the barcode detection of the 2,916 combinations. Sequences of PCR primers used for library amplification and validation are listed. For RNAmap, the sequences of 36 readout probes and their corresponding 319 36-bit codes are listed, as well as the readout scheme and the codebook for the 12 selected transcripts profiled in the DDR364 irradiation screen.

**Supplementary Table 7. Cells passed QC in the irradiation base-editing screening and the optical features.** This table lists 226,369 cells in the irradiation base-editing screening that passed the QC criteria, related to Figure 3 and Supplementary Figure 3. The sgRNA_ID identified for each cell matches the sgRNA_ID in Supplementary Table 5. The average nuclear intensity of protein stains, cell cycle phase annotation, the raw count for foci features, and the number of RNA-reporting spots measured by RNAmap for the 12 transcripts are listed. Cells are then grouped by sgRNA_ID for FDR and LFC calculation listed in Supplementary Table 8.

**Supplementary Table 8. Base-editing screening data analysis.** The KS test p values, FDR, and log2-fold change (LFC) of each guide on each foci features in the irradiation (irradiation) and DNA damaging agents (chemo) screening are shown, the sgRNA_IDs, Rule Set 2 on-target scores, sgRNA categories and Clinvar categories are listed alongside, related to Figure 4, 5 and Supplementary Figure 4, 6. P values of gene enrichment fisher test are listed, related to Figure 4 and Supplementary Figure 3K&N. The wasserstein distances for all foci features are shown, related to Supplementary Figure 4a&B. The p values of beta binomial tests for all binary features are shown, related to Supplementary Figure 4c. The FDR and LFC of foci features for the 9 individually transduced guides are shown, related to Supplementary Figure 4d&E. The number of RNAmap spots and TPM RNA-seq reads for the 12 transcripts are listed, related to Figure 3E.

